# The ClpX and ClpP2 Orthologs of *Chlamydia trachomatis* Perform Discrete and Essential Functions in Organism Growth and Development

**DOI:** 10.1101/868620

**Authors:** Nicholas A. Wood, Amanda M. Blocker, Mohamed A. Seleem, Martin Conda-Sheridan, Derek J. Fisher, Scot P. Ouellette

**Affiliations:** Department of Pathology and Microbiology, College of Medicine, University of Nebraska Medical Center, Omaha, NE; School of Biological Sciences, Southern Illinois University Carbondale, Carbondale, IL; Department of Pharmaceutical Sciences, College of Pharmacy, University of Nebraska Medical Center, Omaha, NE

**Keywords:** *Chlamydia*, differentiation, division, protein turnover, protein quality control, Clp protease, ClpX, ClpP

## Abstract

*Chlamydia trachomatis* (Ctr) is an obligate intracellular bacterium that undergoes a complex developmental cycle in which the bacterium differentiates between two functionally and morphologically distinct forms, the EB and RB, each of which expresses its own specialized repertoire of proteins. Both primary (EB to RB) and secondary (RB to EB) differentiation require protein turnover, and we hypothesize that proteases are critical for mediating differentiation. The Clp protease system is well conserved in bacteria and important for protein turnover. Minimally, the system relies on a serine protease subunit, ClpP, and a AAA+ ATPase, such as ClpX, that recognizes and unfolds substrates for ClpP degradation. In *Chlamydia*, ClpX is encoded within an operon 3’ to *clpP2*. We present evidence that the chlamydial ClpX and ClpP2 orthologs are essential to organism viability and development. We demonstrate here that chlamydial ClpX is a functional ATPase and forms the expected homohexamer *in vitro*. Overexpression of a ClpX mutant lacking ATPase activity had a limited impact on DNA replication or secondary differentiation but, nonetheless, reduced EB viability with observable defects in EB morphology noted. Conversely, the overexpression of a catalytically inactive ClpP2 mutant significantly impacted developmental cycle progression by reducing the overall number of organisms. Blocking *clpP2X* transcription using CRISPR interference led to a decrease in bacterial growth, and this effect was complemented *in trans* by a plasmid copy of *clpP2*. Taken together, our data indicate that ClpX and the associated ClpP2 serve distinct functions in chlamydial developmental cycle progression and differentiation.

**Importance**

*Chlamydia trachomatis* is the leading cause of infectious blindness globally and the most reported bacterial sexually transmitted infection both domestically and internationally. Given the economic burden, the lack of an approved vaccine, and the use of broad-spectrum antibiotics for treatment of infections, an understanding of chlamydial growth and development is critical for the advancement of novel, targeted antibiotics. The Clp proteins comprise an important and conserved protease system in bacteria. Our work highlights the importance of the chlamydial Clp proteins to this clinically important bacterium. Additionally, our study implicates the Clp system playing an integral role in chlamydial developmental cycle progression, which may help establish models of how *Chlamydia* spp. and other bacteria progress through their respective developmental cycles. Our work also contributes to a growing body of Clp-specific research that underscores the importance and versatility of this system throughout bacterial evolution and further validates Clp proteins as drug targets.

## Introduction

*Chlamydia trachomatis* (Ctr) is the leading cause of both bacterial sexually transmitted infections (STIs) and infectious blindness worldwide (1, 2). When left untreated in women, STIs can result in chronic sequelae, including pelvic inflammatory disease, ectopic pregnancy, and tubal infertility. A better understanding of Ctr molecular processes may help reveal essential systems that can be leveraged for more targeted intervention strategies.

*C. trachomatis* is an obligate intracellular bacterial pathogen that differentiates between distinct functional and morphological forms during the course of its developmental cycle (see (3) for review). The elementary body (EB) is small (∼0.3 μm in diameter), infectious, but non-dividing (4, 5). EBs attach to host cells and are internalized into host membrane-derived vacuoles that are rapidly modified into the inclusion (6–9). Within this inclusion, EBs undergo primary differentiation into larger (∼1.0 μm in diameter) reticulate bodies (RBs). RBs are non-infectious but divide using a polarized budding mechanism (10) until secondary differentiation from RBs to EBs occurs. Many studies have detailed the transcriptional and proteomic differences between EBs and RBs (11–14). Given that chlamydial differentiation is not preceded by an unequal division and redistribution of intracellular proteins, as occurs in other bacteria like *Bacillus subtilis* (see (15) for review) or *Caulobacter crescentus* (16), and that EBs and RBs have distinct proteomes, we hypothesize that proteomic turnover plays an integral role in chlamydial differentiation.

Previously, our groups characterized the two ClpP paralogs of *C. trachomatis*. We established that the *clp* protease-associated genes are expressed mid-developmental cycle and that ClpP1 and ClpP2 perform unique functions in chlamydial physiology based on differential effects related to overexpression of catalytically inactive mutants (17). In addition to the two ClpP paralogs, Ctr encodes ClpX and ClpC orthologs (18). ClpX and ClpC are Type I AAA+ (**<u>A</u>**TPase **<u>A</u>**ssociated with diverse cellular **<u>A</u>**ctivities) unfoldases that utilize ATP hydrolysis to linearize protein substrates for either degradation by the ClpP protease or refolding (19, 20). Type I AAA+ ATPases encode Walker A and Walker B motifs, which are responsible for ATP binding and hydrolysis, respectively (21, 22). These AAA+ unfoldases oligomerize to form homo-hexamers that then recognize substrates through multiple different mechanisms (see (23) for review).

Here, we characterized the role of ClpX in chlamydial growth and development. Because ClpX is encoded within an operon with ClpP2, we also investigated effects of overexpression and knockdown of both components. Ctr ClpX is highly conserved, possesses ATPase activity, and formed the expected homohexamer *in vitro*. Interestingly, overexpression of wild-type ClpX, ClpP2, and ClpP2X constructs in Ctr had little effect on bacterial growth, but overexpression of the inactive mutants (alone or in tandem) had a substantial negative effect on recoverable inclusion forming units (IFUs). However, the reduction in IFUs upon inactive ClpX overexpression resulted from non-functional EB generation while the IFU reduction upon inactive ClpP2 overexpression was the result of a block in developmental cycle progression. Our results indicate that chlamydial ClpX is a true ortholog of bacterial ClpX and that the ClpP2X gene products are integral to chlamydial growth and development.

## Results

### The chlamydial ClpX retains conserved motifs of, and exhibits predicted structural homology to, ClpX orthologs

To initiate our study, we first performed bioinformatic and *ab initio* structural modeling analyses to determine whether the chlamydial ClpX (ClpX_Ctr_) possesses the expected conserved regions and motifs consistent with its proposed function as an AAA+ ATPase. Using multiple sequence alignment, we aligned ClpX_Ctr_ to ClpX orthologs and annotated conserved motifs identified in other studies (Fig. 1a). ClpX_Ctr_ retains the N-terminal metal binding domain (24, 25), the Walker A and B motifs for ATP binding and hydrolysis, respectively (21, 23), the sensor motifs for recognition of nucleotide bound state (26), the RKH motif and pore loops for substrate recognition (27–29) and unfolding (30, 31), the arginine finger for inter-subunit sensing of nucleotide state in the ClpX hexamer (22, 32), and the IGF Loop for interaction with ClpP (33, 34). Interestingly, the predicted secondary structure of ClpX_Ctr_ shows few notable aberrations (see Discussion) from other prototypical bacterial ClpX orthologs and is predicted to form the expected homohexamer by structural modeling (Fig. 1b, two subunits removed for clarity). The spatial conservation of AAA+ and ClpX-specific motifs (colored in Fig. 1b as in the multiple sequence alignment) indicates that the chlamydial ClpX likely functions using a mechanism similar or identical to other ClpX orthologs. Taken together, these *in silico* studies suggest that ClpX_Ctr_ functions as a canonical AAA+ ATPase.

**Figure 1:**
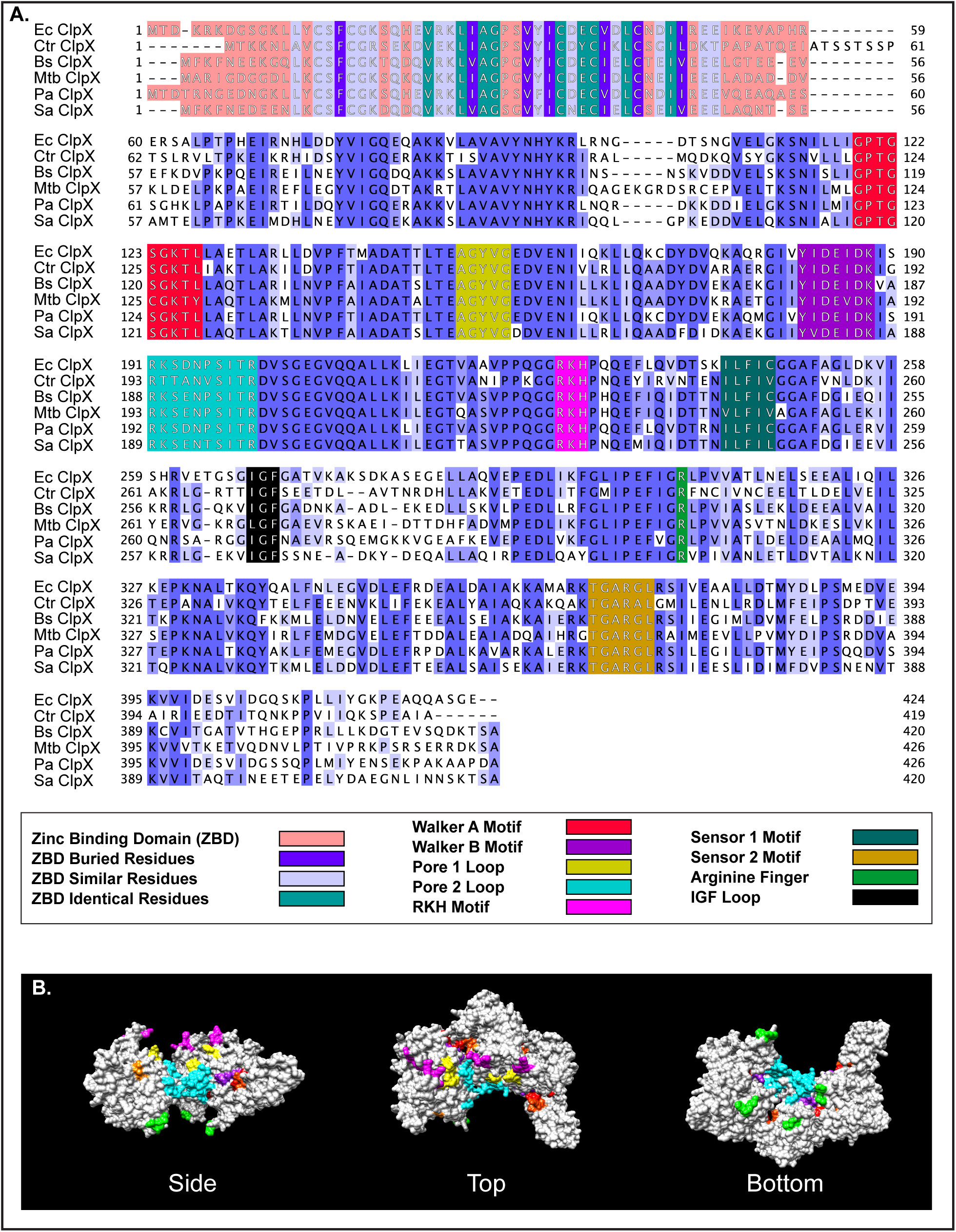
Bioinformatic analysis of chlamydial ClpX supports its role as a AAA+ ATPase. **(A) Multiple sequence alignment** of chlamydial ClpX with the ClpX orthologs of various other bacteria. Ec = *Escherichia coli*, Ctr = *Chlamydia trachomatis*, Bs = *Bacillus subtilis*, Mtb = *Mycobacterium tuberculosis*, Pa = *Pseudomonas aeruginosa*, Sa = *Staphylococcus aureus*. Alignment was performed using Clustal Omega with default settings and presented using Jalview version 2. Alignment was colored by % identity in shades of blue or as indicated below the alignment. (**B) 3D model** of ClpX was generated using SwissModel and presented in UCSF Chimera. Conserved motifs colored as above in (A) except the IGF loops, which are colored lime green. Two subunits of the hexamer were hidden for easier visualization into the complex. Top and bottom are representations following a 90° rotation either clockwise or counterclockwise around the X axis. Of note, this model was generated using an ADP-bound form of ClpX as a template.

### Chlamydial ClpX forms the expected homohexamer and possesses ATPase activity

To determine the oligomeric state of ClpX_Ctr_ *in vitro*, we purified recombinant protein and analyzed its migration by native PAGE. At the same time, we also constructed a Walker B ATPase mutant (E187A) ClpX_Ctr_ as a control for biochemical studies. Following the incubation of 10 μg of wild-type or mutant ClpX_Ctr_ for 20 minutes in a HEPES based buffer, we loaded the entire volume into a 4-20% gradient gel. We observed the ClpX_Ctr_ proteins migrating above the 242 kDa band of the molecular weight ladder, which is close to the expected hexameric size of 283 kDa (Fig. 2a). We then sought to assess ATPase activity of recombinant wild-type and ATPase mutant ClpX_Ctr_ using the Kinase-Glo® endpoint assay, which measures ATP remaining in a sample at the end of the reaction period. Indeed, ClpX_Ctr_ hydrolyzed ATP, while the ATPase mutant isoform showed a significant defect in ATP hydrolysis (Fig. 2b). These data indicate that ClpX_Ctr_ (i) forms a homohexamer of the predicted size and (ii) possesses ATPase activity that is abrogated by a mutation in the Walker B motif.

**Figure 2:**
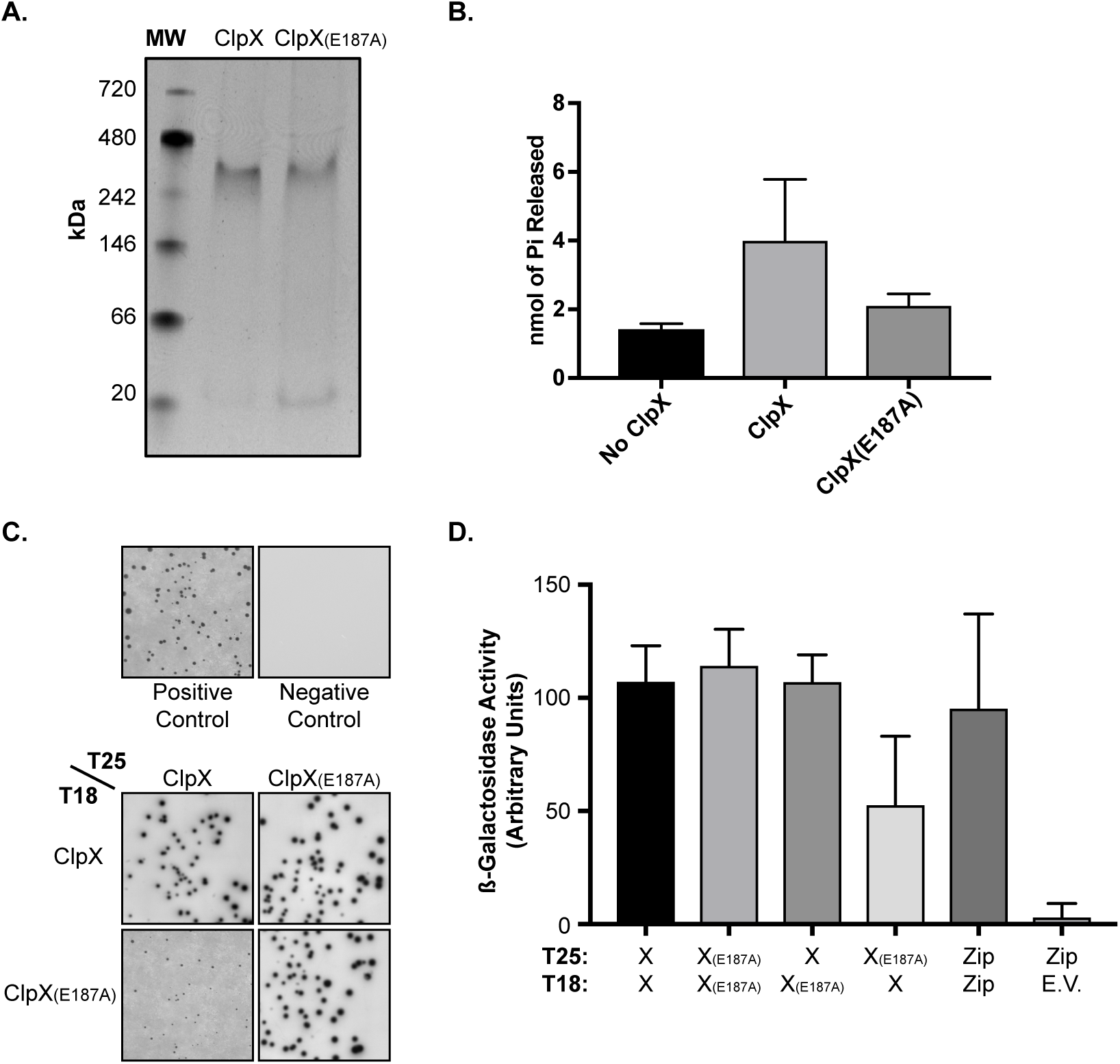
Ctr ClpX is a functional ATPase that forms the expected hexamer. **(A) Native-PAGE assay** of recombinant ClpX and ClpX_(E187A)_. Expected hexameric size is approximately 283 kDa. **(B) ATP hydrolysis** end point assay using Kinase-Glo®. Relative luminescence units (RLU) are displayed on the Y axis. Error bars represent standard deviation, and differences between samples are significant (**** = p < 0.0001, one-way ANOVA). **(C) Bacterial Adenylate Cyclase Two Hybrid (BACTH)** assays showing pairwise, homotypic interaction of ClpX and ClpX_(E187A)_ as well as heterotypic interaction of ClpX and ClpX_(E187A)_. **(D)** β**-Galactosidase activity** of the BACTH interactions from (C), displayed in arbitrary units on the Y axis. All conditions p < 0.05 by unpaired t test compared to negative control.

We next tested whether wild-type and ATPase mutant ClpX_Ctr_ interact with each other using the Bacterial Adenylate Cyclase Two-Hybrid (BACTH) assay. This system is predicated on the reconstitution of adenylate cyclase activity by bringing two complementary fragments of the enzyme (T25 and T18) into close proximity by interacting proteins. Generation of cAMP by the reconstituted adenylate cyclase drives ß-galactosidase production that can be measured qualitatively by the presence of blue colonies and growth on minimal medium (Fig. 2c) or quantitatively by measuring enzyme activity directly (Fig. 2d). We performed a series of pairwise interaction tests between the wild-type and mutant ClpX_Ctr_. In each instance, we observed a positive interaction that was quantifiable and on par with the positive control (T25-Zip vs T18-Zip). We conclude from these data that the mutant isoform can interact with the wild-type isoform.

### Overexpression of ATPase mutant ClpX or catalytically inactive ClpP2 has both distinct and overlapping effects

We previously measured the effects of overexpression of both wild-type and catalytically inactive ClpP2_Ctr_ on chlamydial growth and observed a modest reduction in growth at 24 hours post-infection (hpi) (17). We wanted to more carefully assess growth differences during the chlamydial developmental cycle in the presence of overexpressed wild-type and mutant ClpX_Ctr_ and ClpX_E187A_, ClpP2_Ctr_ and ClpP2_S98A_, or both together (ClpP2X_Ctr_ and ClpP2_S98A_ClpX_E187A_). To do this, we performed growth curves where we induced expression, or not, at 10 hpi and quantified growth at various timepoints after induction. Immunofluorescence analysis (IFA) of replicate treatments and quantification of recoverable inclusion forming units (IFUs; a proxy for EBs) revealed distinct effects upon overexpression of the individual components (Fig. 3a-c) as well as with the entire operon (Fig. 3d&e). We noted that overexpression of wild-type ClpP2_Ctr_ showed no appreciable effect at either 24 or 48 hpi (14 and 38 h pulses of induction, respectively), whereas overexpression of ClpP2S98A appeared to reduce the number of organisms present within the inclusion at 48 hpi but not at 24 hpi (Fig. 3a). These observations correlated with a marked decrease in EB production in the later time points of mutant ClpP2_S98A_ but not wild-type ClpP2_Ctr_ overexpression (Fig. 3b). Conversely, ATPase mutant ClpX_E198A_ overexpression resulted in smaller inclusions and a decrease in IFUs (Fig. 3a&c). Effects on IFU recovery suggest that Ctr is more sensitive to ClpXCtr rather than ClpP2Ctr disruption earlier in the developmental cycle, as the IFU reduction is exacerbated sooner with ClpX_E198A_ overexpression (observe the differences at 24hpi in Fig. 3b&c). As noted for the overexpression of individual wild-type isoforms, there was no significant impact on IFU recovery of overexpressing both wild-type ClpP2_Ctr_ and ClpX_Ctr_ in tandem. Consistent with the effects of overexpressing individual mutant isoforms, overexpression of the mutant ClpP2_S98A_ and ClpX_E187A_ isoforms in tandem showed an exacerbated phenotype throughout the developmental cycle as noted by both IFA and IFU assays (Fig. 3d&e). Importantly, the wild-type chromosomal copies of ClpP2_Ctr_ and ClpX_Ctr_ continue to be expressed during these overexpression assays. Therefore, the true impact of overexpression of the mutant isoforms is likely underrepresented.

**Figure 3:**
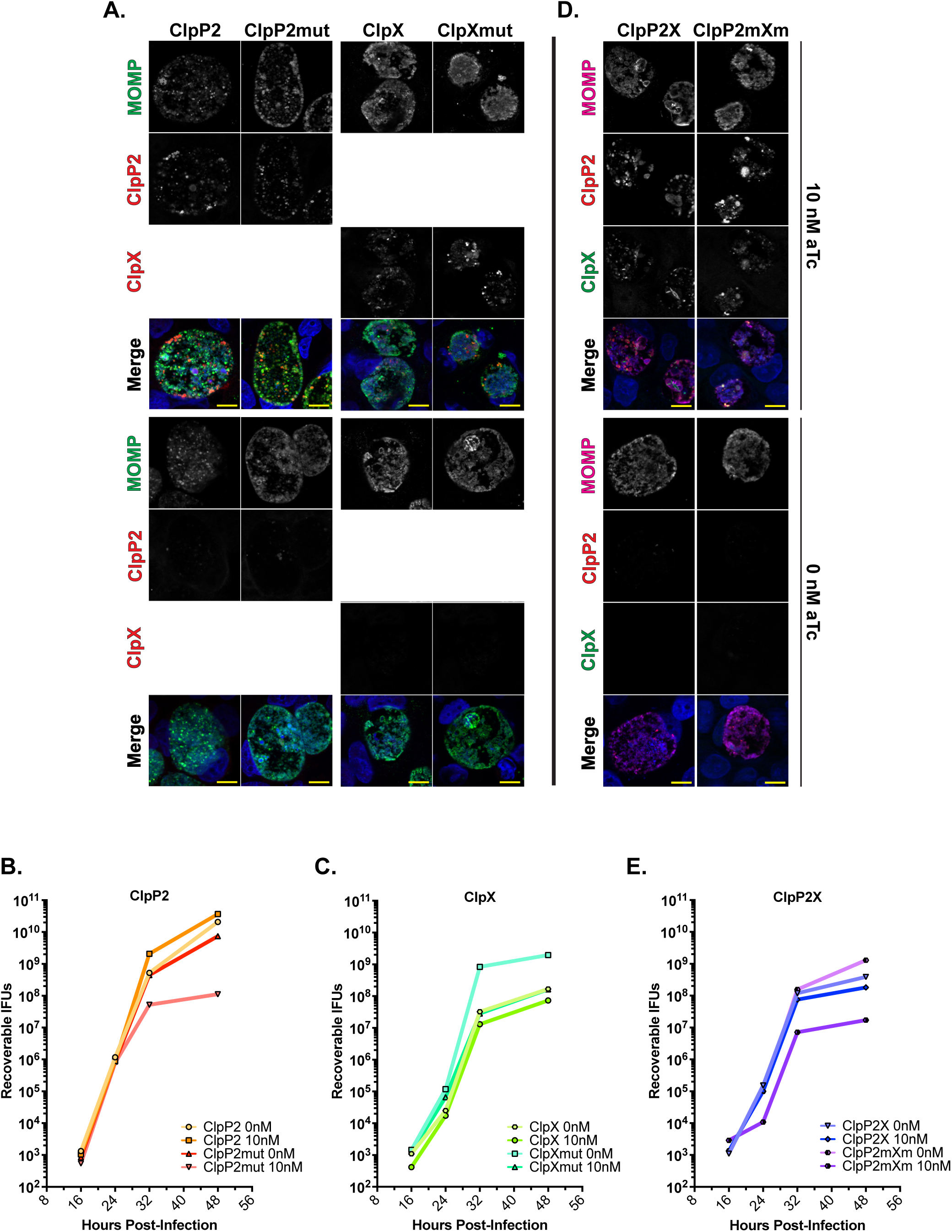
Overexpression of Clp mutant isoforms negatively impacts Ctr at later timepoints. **(A) IFA of ClpP2 and ClpX** wild-type and mutant overexpression at 24 and 48 hpi. Samples were induced with 10 nM aTc at 10 hpi and stained for MOMP (green), 6xHis (red), and DNA (blue). Scale bar = 10 μm. Images were acquired on a Zeiss Apotome at 100x magnification. **(B-C) One step growth curves** of wild-type and mutant ClpP2 (B) or ClpX (C). Samples induced with 10 nM aTc at 10 hpi. IFUs recovered displayed as Log_10_. Values represent the average of two independent experiments. **(D) IFA of ClpP2X** wild-type or mutant operons at 24 and 48 hpi. Samples stained for MOMP (pink), FLAG (ClpP2, red), 6xHis (ClpX, green), and DNA (blue). Parameters as described in (B). **(E) One step growth curves** of ClpP2X wild-type and mutant overexpression. IFUs recovered displayed as Log_10_.

### Functional disruption of ClpP2 blocks developmental cycle progression while ClpX disruption reduces EB viability

Given that the IFU assay only measures EB viability from a population and not total bacterial numbers or differentiation status, we next wanted to address these nuances of the chlamydial developmental cycle. We first measured genomic DNA as a proxy for total number of bacteria (i.e. both RBs and EBs). Overexpression of any wild-type protein had no significant impact on DNA accumulation. From 24 hpi to 48 hpi, we observed a significant drop in the rate of increase of gDNA levels when ClpP2_S98A_ was overexpressed alone or in the mutant operon configuration (Fig. 4a). Overexpression of the ClpX_E187A_ showed a trend (two-fold) towards decreased DNA levels that was not statistically significant. This was surprising given the roughly 20-fold reduction in IFUs (Fig. 3), suggesting total bacterial numbers during ClpX_E187A_ overexpression are unaffected.

**Figure 4:**
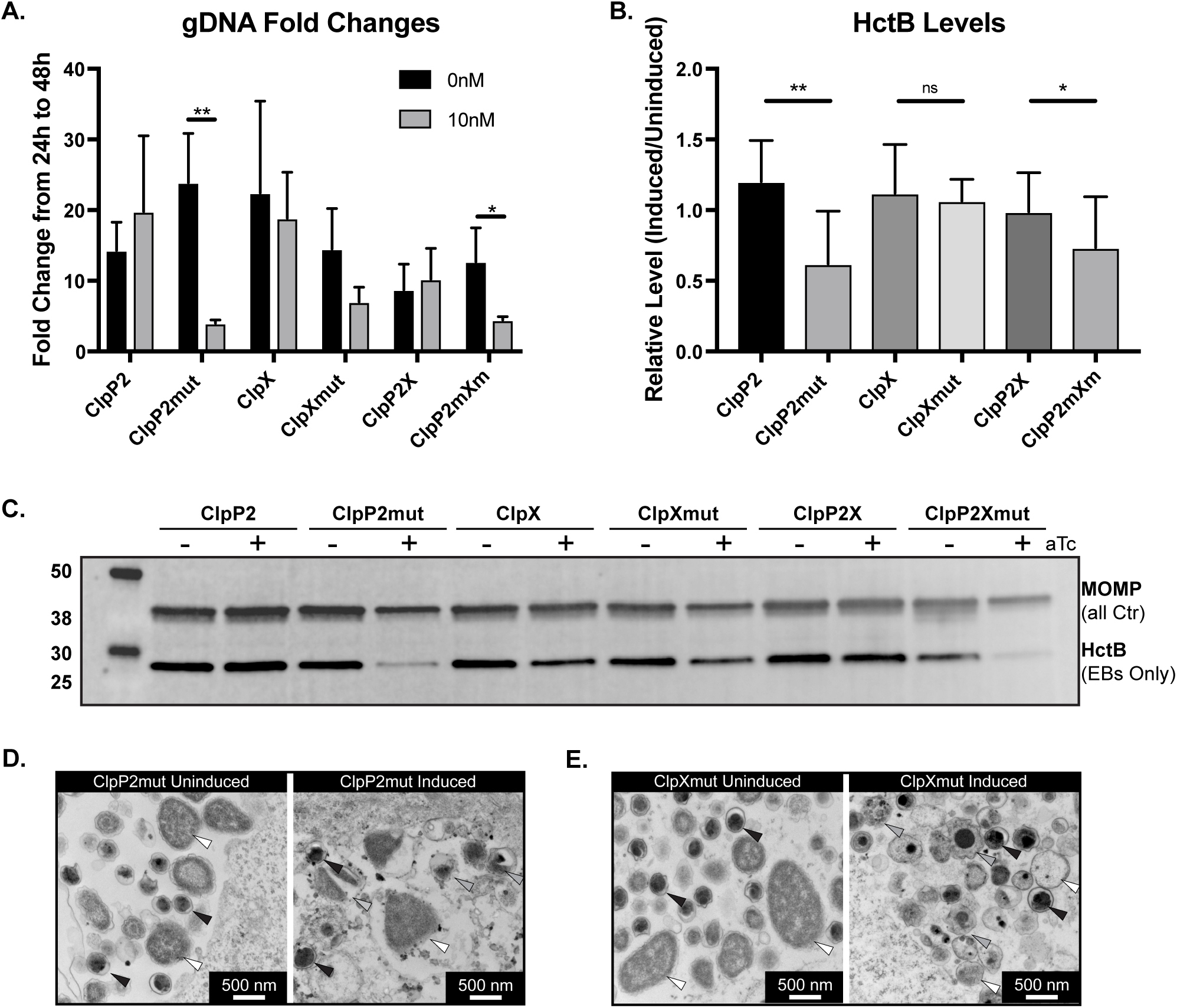
Functional disruption of ClpX and ClpP2 perturbs chlamydial development but in different manners. **(A) Fold changes** of detectable gDNA from 24 to 48 hpi for each strain. Samples were induced at 10 hpi with 10 nM aTc. * = p ≤ 0.05, ** = p ≤ 0.001 by paired t-test. Values displayed as the average of three independent experiments with error bars representing standard deviation. **(B) Western blot** analysis of HctB levels at 48 hpi with and without overexpression. One well of a six well plate was lysed into 500 μL of denaturing lysis buffer. 50 μg of protein from the clarified lysate for each sample was assessed. Blots were probed for MOMP (IR680) and HctB (IR800). Grayscale blot shown is representative of three independent experiments. **(C) Quantified integrated** density of the staining from (B). HctB levels were normalized to MOMP levels in each sample to account for differences in bacteria in each sample. Values displayed as levels of induced to uninduced HctB/MOMP ratios for each strain. * = p ≤ 0.05, ** = p ≤ 0.001, ns = not significant by multiple comparisons t-test. **(D-E) Representative images** of electron micrographs of the indicated strains and conditions. White triangles indicate normal RBs, black triangles indicate normal EBs, and gray triangles indicate abnormal developmental forms.

To determine if there was an impact on differentiation status during overexpression of wild-type and mutant isoforms, we next assessed HctB levels, an EB-specific gene product (35–37), by western blot as an indicator of secondary differentiation. We normalized the integrated density of HctB to the integrated density of MOMP (major outer membrane protein; present in both EBs and RBs) to ensure that we were comparing HctB levels to the total number of bacteria to give a more robust readout of differentiated bacteria within the population. The relative HctB levels in samples where ClpP2_S98A_ was overexpressed were significantly reduced as expected, indicating the generation of fewer EBs and consistent with IFU and genomic DNA data, whereas the other experimental conditions showed no changes in relative HctB levels (Fig. 4b&c). These data suggest that overexpression of ClpX_E187A_ does not impact total number of bacteria, as measured by gDNA levels, or RB-to-EB differentiation, as measured by HctB levels. Nevertheless, the recoverable IFUs were decreased when ClpX_E187A_ is overexpressed (Fig. 3C&E), indicating a defect in the EBs being produced.

To further explore this possibility, we prepared samples for transmission electron microscopy to examine at higher resolution the morphology of EBs and RBs from ClpP2_S98A_ and ClpX_E187A_ overexpressing strains. As RBs condense to EBs, an intermediate form can be observed where the chlamydial nucleic acids are visible as an electron dense spot. Consistent with other measured effects (Fig. 4d&e; Suppl. Fig. S1a), ClpP2_S98A_ overexpression resulted in smaller inclusions with fewer organisms whereas ClpX_E187A_ appeared to have no effect on bacterial numbers (Fig. 4d&e). In both induced and uninduced samples for each strain, we noted normal RBs (Fig. 4d&e, white triangles) and normal EBs (Fig. 4d&e, black triangles). However, we observed numerous abnormal developmental forms following ClpX_E187A_ overexpression that appeared to be multi-nucleated or abnormally condensing intermediate forms (Fig. 4e, gray triangles). These data are consistent with the limited effect of ClpX_E187A_ overexpression on the production of HctB (Fig. 4b&c) but the decreased recoverable IFUs (Fig. 3a&c), further supporting that disruption of normal ClpX function may affect EB viability.

### Knockdown of the clpP2X operon reduces recoverable progeny and results in reduced plasmid retention while complementation with clpP2 partially restores infectivity

Overexpression of mutant isoforms of ClpP2_Ctr_ and/or ClpX_Ctr_ was sufficient to disrupt chlamydial development in the presence of endogenous ClpP2X_Ctr_. However, we wanted to directly block the chromosomal copies by employing an improved version of the chlamydial CRISPR interference (CRISPRi) strategy previously described by us ((38) and Ouellette et al, *in prep*). CRISPRi relies on the inducible expression of a catalytically inactive Cas9 (dCas9) in combination with a guide RNA (gRNA) to block transcription at specific chromosomal sites (39). We transformed Ctr L2 with vectors encoding inducible dCas9 and constitutive gRNAs targeting either the *clpP2X* or *incA* intergenic regions (Fig. 5A and Suppl. Figs. S2&3). Of note, *pcnB2*, a predicted poly-A polymerase, is encoded at the 3’ end of the *clpP2X* operon and was included in our analyses. To determine whether we could also complement the effects of knockdown, we constructed vectors encoding either *clpP2_FLAG* or *pcnB2_FLAG* in a transcriptional fusion 3’ to dCas9. These vectors were transformed into *C. trachomatis* L2. Under conditions when dCas9 is expressed, ClpP2_FLAG or PcnB2_FLAG is also expressed (Fig. 5b and Suppl. Fig. S4). IncA knockdown served as a control since *incA* is a non-essential gene (40). These CRISPRi transformants were used to infect HEp2 cells.

**Figure 5:**
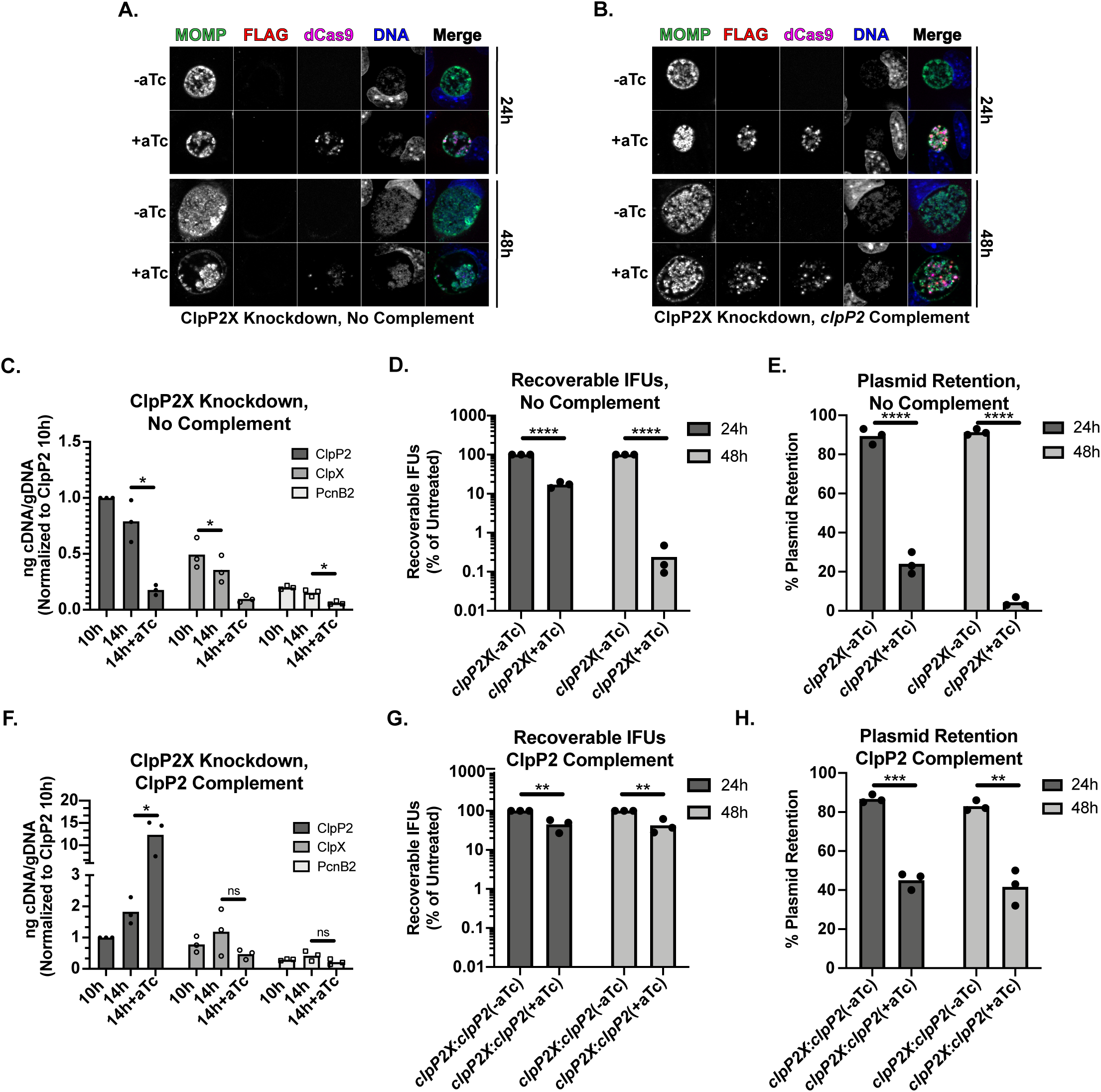
Knockdown of ClpP2X negatively impacts *Chlamydia*, while complementation with ClpP2 partially restores normal development. **(A-B) Representative immunofluorescence** images of *clpP2X* knockdown with either (A) no complementation or (B) *clpP2* complementation. **(C) Transcript levels** upon knockdown of *clpP2X*. Data displayed are the values of three independent biological replicates of triplicate RT-qPCR reactions. Values are normalized to the 10h uninduced ClpP2 value for each experiment. Samples were induced using 10 nM aTc. **(D) IFU titer** following induction of knockdown at 4 hpi. Values are presented on a Log_10_ scale percent of the respective uninduced titer. **(E) Plasmid retention** based on the ratio of GFP positive to total number of inclusions is displayed in percent for each condition. **(F-H) IFU titer and plasmid retention** of *clpP2X* knockdown with *clpP2* complementation. Conditions used are the same as in (C-E).

When dCas9 expression was induced at 10hpi in the various transformants, we observed a marked and rapid decrease in *clpP2*, *clpX*, and *pcnB2* transcript levels compared to the uninduced controls at 14hpi (Fig. 5c). Importantly, we did not observe a decrease in transcript levels for unrelated genes associated with different stages of the developmental cycle (*clpP1, euo,* and *omcB*; Suppl. Fig. S2; (12, 17, 41, 42)). We also observed abnormal inclusion morphology with individual organisms aggregated to one side of the inclusion (Fig. 5a). Complementation with ClpP2_FLAG resulted in decreased *clpX* transcripts as compared to the uncomplemented strain but an increased level of *clpP2* transcripts, as expected (Fig. 5f). However, ClpP2 complementation did not fully restore inclusion morphology as organisms still displayed some aggregation within the inclusion, albeit intermediate between the uncomplemented knockdown and control conditions (Fig. 5b). Complementation with PcnB2_FLAG did not impact *clpP2* and *clpX* transcript levels during knockdown but did increase *pcnB2* transcripts approximately 10-fold relative to the uninduced samples (Suppl. Fig. S4b). Abnormal inclusion morphology was also evident (Suppl. Fig. S4a). As previously observed, IncA expression was uniformly blocked after dCas9 induction (Suppl. Fig. S3; (38)). We also attempted a similar *clpX* complementation strategy but saw no effect on *clpP2* or *pcnB2* transcript levels during knockdown (unpublished observation). Given that we tagged our dCas9 variant with an SsrA recognition motif and that ClpX recognizes the SsrA tag (see below), the dCas9 was likely targeted for degradation by the plasmid-encoded ClpX, subsequently eliminating any knockdown.

We next assayed chlamydial growth as measured by IFU recovery and chlamydial morphology by immunofluorescence after inducible knockdown of the target genes. Expression of dCas9 was induced at 4hpi, and IFUs were harvested at 24 and 48hpi and titred on fresh cell monolayers in the presence of penicillin, the selection agent. When *clpP2X* expression was blocked at 4hpi, we noted a 5-fold decrease in penicillin-resistant (i.e. transformants containing the CRISPRi plasmid) IFUs at 24hpi but a more than 200-fold decrease at 48hpi (Fig. 5d). Immunofluorescence assay images supported this drop in viable organisms, revealing a severely diminished number of organisms within the inclusion (Fig. 5a). In performing these assays in the presence of penicillin, we also observed numerous penicillin-sensitive organisms (i.e. aberrant RBs (43)) during the titration step, suggesting that the plasmid conferring resistance and encoding the CRISPRi system was being lost after induction of dCas9 expression. To test this, we quantified plasmid retention in the *clpP2X* knocked down samples and observed that blocking *clpP2X* expression resulted in ∼75% plasmid loss at 24hpi and greater than 90% loss at 48hpi (Fig. 5e), which was closely recapitulated with *pcnB2* complementation (Suppl. Fig. S4c&d). These effects on IFUs and plasmid retention were not observed for *incA* knockdown (Suppl. Fig. S3b&c). We note that *incA* knockdown did result in a reproducible, but transient, increase in IFUs at 24hpi that returned to “normal” levels at 48hpi (Fig. S3b). The reasons for this are not clear. Nonetheless, we conclude from these data that blocking *clpP2X* expression is deleterious to *Chlamydia*, further highlighting its essentiality to this pathogen. Interestingly, complementation with *clpP2* largely rescued the drop in recoverable IFUs while maintaining plasmid loss as compared to the uncomplemented condition (Fig. 5g&h). This suggests that much of the loss in bacterial viability during *clpP2X* knockdown is attributable to reduced ClpP2 levels.

### ClpX function is required for degradation of an SsrA-tagged substrate in Ctr

Recently, ClpX-specific inhibitors were synthesized by the Sieber group and were shown to interfere with ClpX ATPase activity (44). One compound, identified as 334, was shown to have potent anti-ClpX activity whereas a derivative, 365, was inactive. We performed *ab initio* modelling and molecular dynamics simulations (45) to determine if these compounds could interact with an ADP-bound hexameric ClpX_Ctr_. For 334, a high scoring model (-9.1 kcal/mol binding affinity, RMSD ∼ 0) was predicted with the drug binding near to the ATP binding pocket, suggesting a mechanism of action where 334 likely occludes the ATPase site (Suppl. Fig. S5). Whether the effect stems from the blocking of ATP binding and subsequent destabilization of the complex, attenuation of ATPase function by preventing a conformational change of the complex, or steric hindrance of complex formation remains to be elucidated. Conversely, compound 365 bound outside of the ATP pocket with a much lower score (Suppl. Fig. S6).

Given the predicted binding of the ClpX inhibitors on the structure of ClpX_Ctr_, we sought to determine whether these drugs effectively blocked ClpX function *in vivo*. We utilized a chlamydial SsrA-tagged GFP variant [GFP(VAA)], which has been previously shown to be a substrate of ClpX_Ctr_ *in vitro* (46), or GFP tagged with a mutated SsrA tag [GFP(VDD)]. In mutating VAA to VDD, GFP is rendered resistant to ClpX recognition (47). Using an anhydrotetracycline-inducible pBOMB4 derivative encoding constitutive mCherry as a metric of chlamydial growth (herein referred to as pBOMBmC), we cloned each GFP variant into pBOMBmC and transformed these constructs into Ctr L2. Samples were infected, and GFP expression was induced at 8 hpi. All induced samples were then allowed to develop until 16 hpi, when media was replaced by either DMEM only or DMEM containing the indicated antibiotic (Fig. 6a&b, Suppl. Fig. S7). Immediately following drug addition, induction of GFP expression was maintained or not, where removal of aTc is denoted by Pulse/Chase. GFP signal was normalized to the constitutive mCherry signal expressed on the plasmid backbone. We observed that removal of inducer results in rapid loss of GFP(VAA) but not GFP(VDD) signal compared to continuous induction, indicating that GFP(VAA) is degraded as expected (Fig. 6a&b, Suppl. Fig. S7). Treatment with compound 334 resulted in sustained GFP(VAA) signal, consistent with its documented effects on ClpX (44). Importantly, compound 365, the inactive 334 derivative, and ACP1b, a ClpXP uncoupling drug with a mechanism of action similar to the ADEP compounds (17, 48), failed to protect GFP(VAA) from degradation (Fig. 6a), which further supports the specificity of compound 334 in mediating this effect. We then tested the effect of these drugs on chlamydial viability and noted a multiple log_10_ decrease in recoverable IFUs (Supp. Fig. S8), supporting that blocking ClpX function is highly detrimental to *Chlamydia*. However, in attempting to assess the effect of these compounds on *Chlamydia*, we noted reduced host cell viability with prolonged treatment (Supp. Fig. S8d), making interpretation of those data unclear.

**Figure 6:**
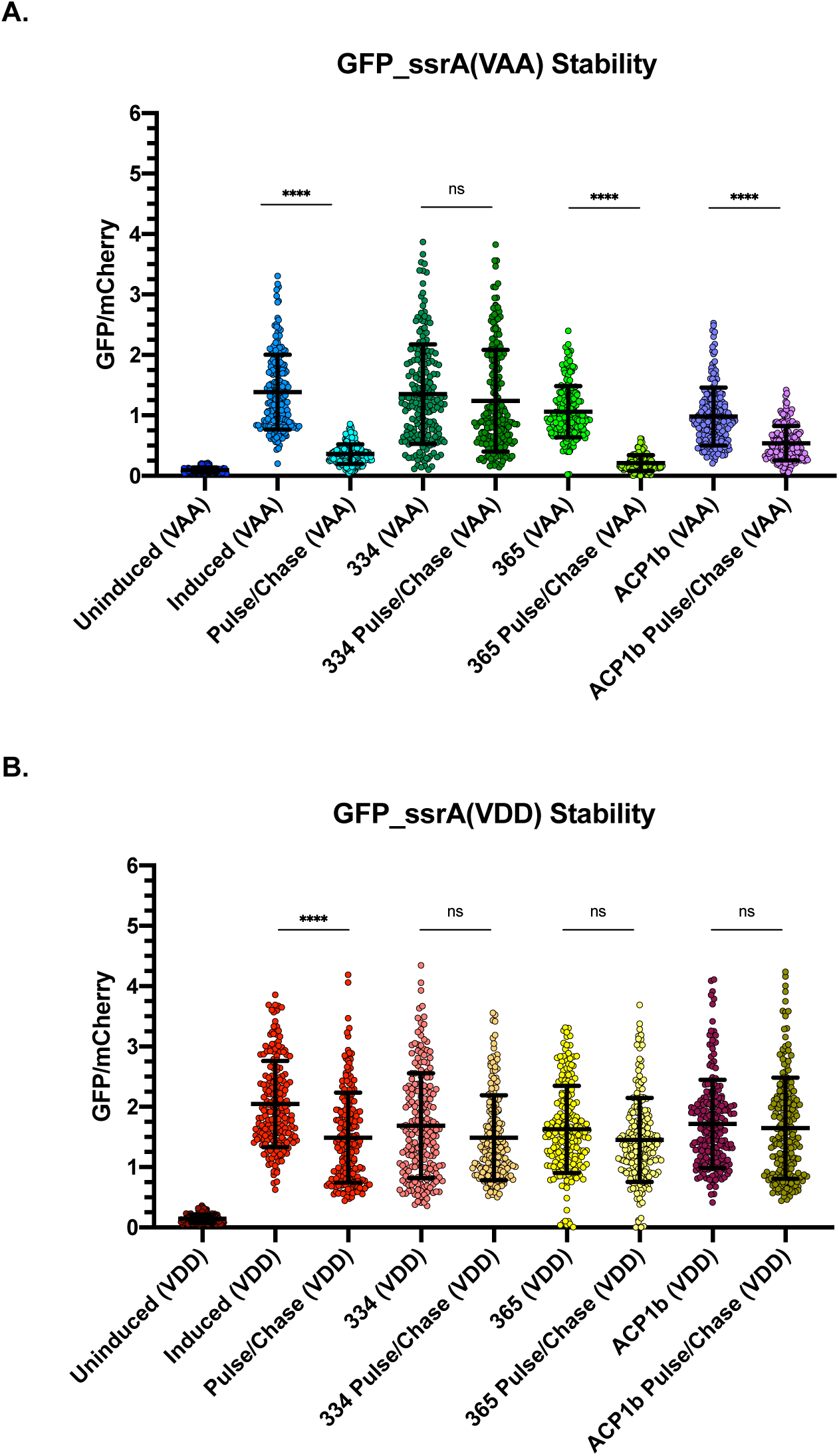
Chemical disruption of ClpX function protects SsrA-tagged GFP. **(A) The ratio** of integrated density measurements of GFP to mCherry for SsrA(VAA)-tagged GFP. All samples were induced or not at 8 hpi with 20 nM aTc. Media was replaced with or without drug at 16 hpi, and aTc induction was maintained or removed (for pulse/chase samples). Samples were fixed at 20 hpi, and GFP and mCherry intensity were quantified using Fiji. **** = p ≤ 0.0001, ns = not significant b**y** ordinary one-way ANOVA. **(B) Same conditions** as in (A) but with GFP(VDD).

## Discussion

Given the unique roles and protein repertoires of the chlamydial developmental forms (EB/RB), we hypothesize that protein degradation is a critical factor in the differentiation process from one form to the other. The Clp system is highly conserved in both prokaryotic and eukaryotic systems where it has been described to perform important functions in both proteostasis and pathogenesis (49). The Clp system is nominally composed of a proteolytic subunit, ClpP, and a AAA+ ATPase that functions as an unfoldase to recognize substrates and feed them into the ClpP barrel for degradation (23). The work presented here expands our understanding of the Clp protease system of an obligate intracellular bacterial pathogen, *Chlamydia trachomatis*. Focusing on characterization of ClpX_Ctr_ and the function of the *clpP2X* operon, we demonstrated the importance of the ClpXP protease system during chlamydial growth and development.

Multiple lines of evidence support that the chlamydial ClpX is a *bona fide* AAA+ ATPase. Firstly, multiple sequence alignment of ClpX_Ctr_ to orthologs of other bacteria revealed a perfect conservation of the motifs involved in nucleotide binding, ATP hydrolysis, and nucleotide-state sensing (Fig. 1) (50, 51). Secondly, homology-directed and *ab initio* modelling of ClpX_Ctr_ revealed that the spatial orientation of these domains is conserved as well (Fig. 1), though we acknowledge that structural studies are critical to drawing conclusions about ClpX_Ctr_ conformational states. Thirdly, ClpX_Ctr_ interacts with itself to form a homohexamer that possesses ATPase activity (Fig. 2). Importantly, this ATPase activity could be disrupted by a targeted mutation in the Walker B motif while having no effect on the oligomerization properties of the protein. Fourthly, a characterized ClpX inhibitor that disrupts its ATPase activity also blocked SsrA-mediated degradation *in vivo* and disrupted the growth of *C. trachomatis* serovar L2 (Fig. 6; Suppl. Fig S7). Finally, overexpression of a ClpX_Ctr_ ATPase mutant *in vivo* negatively impacted chlamydial growth and development (Figs. 3&4).

While we have characterized the ATPase function of ClpX_Ctr_ and its role in chlamydial growth, further work remains to determine whether this ClpX ortholog functions as an unfoldase. Nevertheless, our bioinformatics analysis supports this as ClpX_Ctr_ retains substrate recognition motifs, including both pore loops and the RKH motif for gripping and translocation of substrates (27–31). *Chlamydia* spp. also encode the tmRNA/ssrA tagging system for ribosomal rescue (18, 52–55), which fits a model where ClpX_Ctr_ may play an integral role in turnover of tagged, partially translated peptides. A recent article, using an *in vitro* assay for SsrA-tagged GFP degradation, suggests this function of chlamydial ClpX may be conserved (46). Our *in vivo* data exhibiting SsrA-tagged GFP protection with the addition of a ClpX inhibitor also suggests ClpX_Ctr_ can target SsrA-tagged substrates, but whether this tagging is for ribosomal rescue or more specific purposes (56, 57) remains to be determined and is currently under investigation by our research group.

In *Chlamydia*, *clpX* is encoded in an operon with *clpP2*. Our data indicate that, not surprisingly, the ClpP2X_Ctr_ system is highly regulated and essential. We previously demonstrated that treatment with ClpP-targeting antibiotics severely impacts growth of *Chlamydia* (17). Here, we performed a systematic analysis of the effects of overexpression of wild-type or inactivated ClpP2X_Ctr_ components. The overexpression of wild-type ClpP2_Ctr_ and/or ClpX_Ctr_ had no biologically or statistically significant effect on chlamydial growth that we could measure.

However, extended overexpression of mutant ClpP2_Ctr_(S98A) and/or ClpX_Ctr_(E187A) resulted in abrogation of chlamydial growth as measured by recovery of infectious progeny. Three observations should be noted. Firstly, the effect of inducibly-expressed proteins is measured in the presence of the endogenous chromosomally-expressed copies of these proteins. Therefore, it is likely that the mutants would have even more dramatic effects on chlamydial growth and that disruption of these components results in lethality. For ClpX_Ctr_, this is supported by the effects of the ClpX inhibitor on *Chlamydia* (Supp. Fig. 8), which effectively stopped chlamydial growth. Secondly, we demonstrated that the mutant proteins could interact *in vitro* with wild-type isoforms (Fig. 2 and (17)). Therefore, we can infer that overexpression of the mutant proteins leads to their incorporation into the endogenous Clp machinery to disrupt or impair its function. Thirdly, to our knowledge, ours is the first study to ectopically express two different tagged proteins in *Chlamydia*, showing both the feasibility of this approach and its potential utility to dissect chlamydial biology.

The overexpression of the catalytically inactive mutant Clp proteins in *Chlamydia* revealed potentially subtle differences in the role of each component in chlamydial growth and development. Surprisingly, we noted a roughly 50% reduction in detectable genomes (Fig. 4A) when ClpX_Ctr_(E187A) was expressed whereas IFUs were reduced roughly 20-fold (Fig. 3). The production of EBs as measured by HctB levels did not appreciably change (Fig. 4b&c). This suggests that, while development is hindered, the drop in IFUs may be due to defective EB viability, infectivity, or inclusion establishment and not a reduction in secondary differentiation *per se*. Support for this comes from electron microscopy images, which revealed unusual morphologies after overexpression of the mutant ClpX_Ctr_ isoform (Fig. 4d; Suppl. Fig. S1). As noted above, the SsrA-mediated degradation of partially translated products may be critical during the differentiation process. As RBs condense to EBs, DNA replication, transcription, and translation are significantly abrogated, and we hypothesize that those partially translated proteins trapped during this differentiation step require degradation. This may explain the effects we observed on decreased EB viability during overexpression of the ClpX mutant isoform. Whether the degradation of such partial peptides occurs during RB-to-EB differentiation or during the *next* infection when an EB differentiates to an RB remains to be investigated.

Conversely, for ClpP2_Ctr_(S98A) overexpression, the substantial IFU decrease coupled with a sharp drop in gDNA levels indicate that ClpP2_Ctr_ plays a role in developmental cycle progression. HctB levels are also significantly reduced, which is consistent with the lack of EB generation. Taken together, these data may indicate that ClpP2_Ctr_ is integral to developmental cycle progression or differentiation and that its function is tightly regulated. We cannot, however, conclude that secondary differentiation is directly affected due to the fact that total organism numbers are severely reduced. Rather, our proposed model suggests that ClpP2_Ctr_ disruption may affect both factors by a mechanism that we are currently working to identify. Conversely, ClpX_Ctr_ may serve a more prominent ClpP2_Ctr_-independent function in differentiation of the organism (Figure 7). Interestingly, several studies on the periplasmic protease, HtrA, in *Chlamydia* have implicated this protease in membrane reorganization (58–60). Therefore, one model may be that the ClpXP system may be required for cytosolic turnover whereas HtrA may be required for membrane reorganization during differentiation.

**Figure 7:**
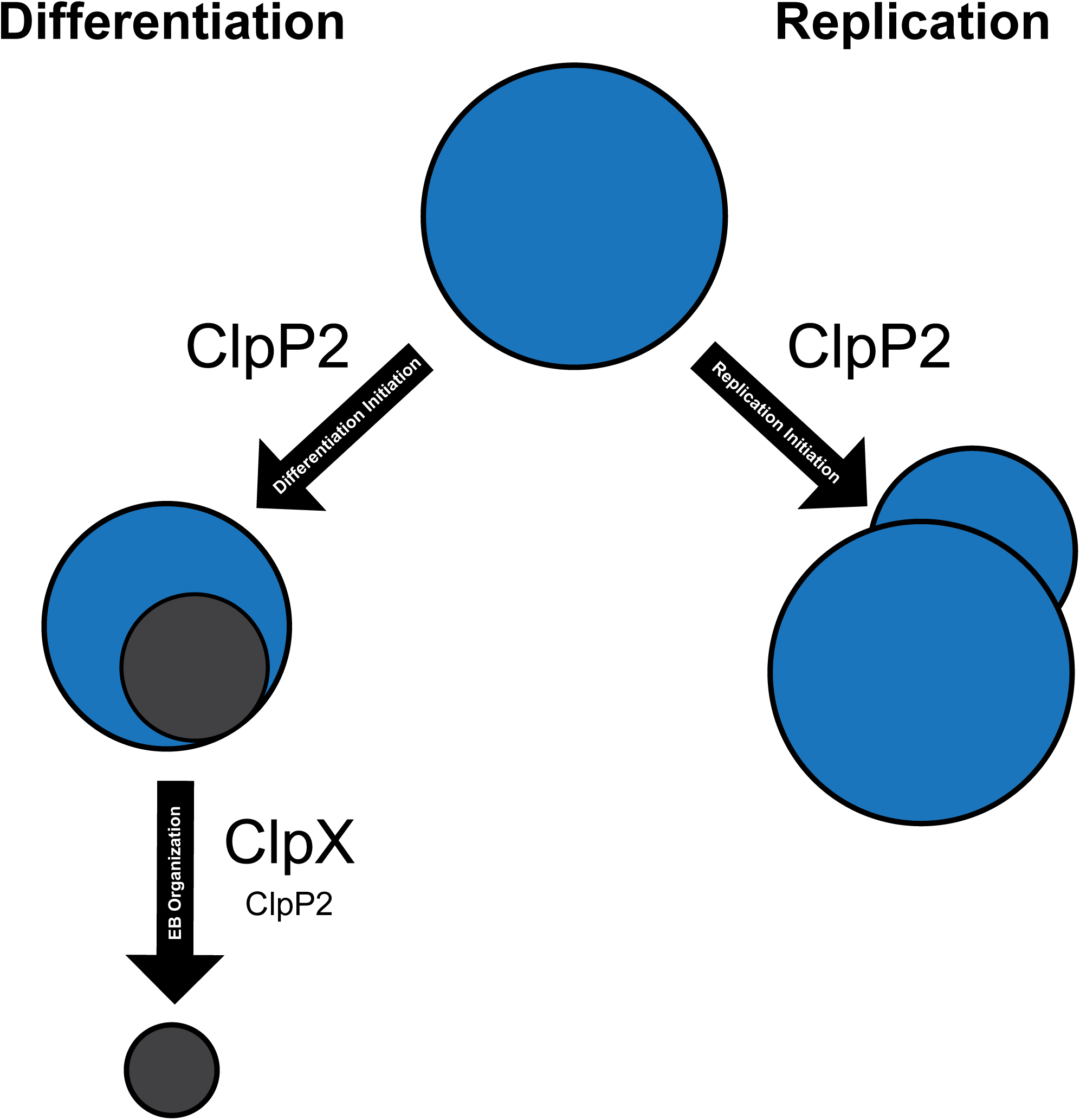
Proposed model for ClpP2X function in *Chlamydia*. An RB ultimately has two fates: differentiation or replication. Based on our data showing the impact of ClpP2 disruption on the developmental cycle, we hypothesize that ClpP2 may function in triggering either event based on its degradative target at either timepoint. Furthermore, we posit that ClpX serves an important function in EB organization, given that ClpX affects recoverable IFUs without reducing the amount of HctB produced.

We successfully generated chlamydial transformants with an inducible knockdown system to repress ClpP2X_Ctr_ expression. To date, this study is the first of its kind in *Chlamydia* to knock down genes that are essential, highlighting the utility of CRISPRi in studies of chlamydial biology while providing insight into possible ClpP2X_Ctr_ function. We also demonstrated that these knockdowns may be complemented by insertion of the gene of interest under the inducible promoter as a transcriptional fusion with the dCas9 (Fig. 5 and Ouellette et al, *in prep*). Notably, we observed a large decrease in IFU production coupled with an increase in plasmid loss after inhibition of *clpP2X* expression (Fig. 5). These effects were not observed when targeting a non-essential gene. Additionally, we demonstrated that the impact on IFUs, but not the effect on plasmid retention, was reversed by complementation. Of note, penicillin does not kill chlamydiae but blocks cell division (61, 62), which keeps the organism transcriptionally in an RB-like state (63). This suggests that knocking down an essential gene(s) puts selective pressure on the chlamydiae to lose the plasmid encoding the CRISPRi system. This has important ramifications for long-term experiments and functional analyses. Nevertheless, the CRISPRi system represents a significant advance for our ability to study essential systems in this obligate intracellular bacterium.

A recent *in vitro* study suggested that homoheptamers of ClpP1 form a complex with homoheptamers of ClpP2 that interact with ClpX (46), yet we observed distinct differences when overexpressing mutant isoforms of these proteins in *Chlamydia* (17). If these ClpP proteins also interact with the other chlamydial AAA+ ATPase, ClpC, then it is not surprising that we observed differences between ClpP2 and ClpX in our assays since disrupting ClpP2 could also be impacting ClpC function. This would also beg the question why we did not observe similar impacts on chlamydial growth when overexpressing each mutant isoform (17). One possibility may be that ClpX interacts with ClpP2 in a ClpP1P2 complex whereas ClpC interacts with ClpP1 in a ClpP1P2 complex. In this scenario, disrupting ClpP2 would act epistatically to ClpX. This is not inconsistent with our data and proposed model. Clearly, the composition of the Clp complexes remains to be determined *in vivo*, and this is an area of ongoing investigation.

In conclusion, we have demonstrated the importance of the ClpP2X_Ctr_ system to chlamydial development, but many questions remain unanswered. These include why ClpP2_Ctr_ and ClpX_Ctr_ may serve independent purposes and what substrates this system may be targeting. Additionally, we need to identify any cofactors, chaperones, adaptor proteins, or a lack thereof that may be pertinent to this system. We plan to dissect the structural motifs of ClpP2_Ctr_ and ClpX_Ctr_ to determine if any of the noted differences from other bacterial Clp proteins may alter activity, which may aid in our goal of further functional assessment. Finally, continued experimentation to address our overarching hypothesis that protein turnover is critical to chlamydial differentiation, and that the Clp system is a key mediator of this process, is required. Overall, we conclude that the chlamydial ClpP2X_Ctr_ system is critical to the development of these obligate intracellular bacteria.

## Materials and Methods

### Strains and Cell Culture

The human epithelial cell line HEp2 was used for the overexpression assays, gDNA and protein extractions, and antibiotic studies. McCoy mouse fibroblasts were used for chlamydial transformation, and human epithelial HeLa cells were used for plaque purification. All of these cell lines were passaged routinely in Dulbecco’s Modified Eagle’s Medium (DMEM, Gibco/ThermoFisher) and 10% FBS (Sigma; St. Louis, MO) and verified to be mycoplasma-free using LookOut^®^ Mycoplasma PCR Detection Kit (Sigma). Density gradient purified *Chlamydia trachomatis* L2/434/Bu (ATCC VR902B) EBs were used for the antibiotic studies. *C. trachomatis* serovar L2 EBs (25667R) naturally lacking the endogenous plasmid were prepared and used for transformation [see (64)].

### Bioinformatics Analysis

Gene sequences of *Chlamydia trachomatis* were obtained from STDGen database (http://stdgen.northwestern.edu) or KEGG Genome Browser (65–67). RefSeq protein sequences from *Escherichia coli, Bacillus subtilis, Mycobacterium tuberculosis, Staphylococcus aureus,* and *Pseudomonas aeruginosa* were acquired from the NCBI protein database (https://www.ncbi.nlm.nih.gov/guide/proteins/). ClpX pairwise protein alignments to find sequence identity were performed using NCBI Protein BLAST function (https://blast.ncbi.nlm.nih.gov/Blast.cgi) (68). Multiple sequence alignments were performed using Clustal Omega (69) with default settings and were presented using Jalview Version 2 (70). PDB files for predicted monomeric 3D structures were acquired from the Phyre2 website (http://www.sbg.bio.ic.ac.uk/phyre2/html/page.cgi?id=index) (71). Complexes were modelled using SWISS-MODEL available on the ExPASy server (72–75). Protein models and model alignments were rendered using the UCSF Chimera package from the Computer Graphics Laboratory, University of California, San Francisco (supported by NIH P41 RR-01081) (76). Docking analyses were performed using AutoDock Vina (45). Molecules were prepped using Dunbrack rotamer libraries (77, 78) to replace incomplete side chains and ANTECHAMBER for charge assignment and topology generation (79).

### Plasmid Construction

A full list of the primers and plasmids used is included in the supplementary material. The Gateway® recombination system of cloning was used for plasmids for the Bacterial Adenylate Cyclase Two-Hybrid (BACTH) system (80). The genes were amplified from *Chlamydia trachomatis* L2 genomic DNA with added *attB* recombination sites. The PCR products were then incubated with a pDONR^TM^221 entry vector (containing *attP* recombination sites) in the presence of BP Clonase II (Invitrogen) to insert the gene via flanking *attP* recombination sites and remove the *ccdB* insert, resulting in an entry vector containing the gene of interest flanked by *attL* sites. These constructs were transformed into DH5α chemically competent *E. coli* and plated onto kanamycin-containing LB agar. Plasmid was isolated and used for the LR reaction into one of three destination vectors (pST25-DEST, pSNT25-DEST, or pUT18C-DEST). The same entry vector for any given gene was used for all three LR reactions to insert into the destination vector. Entry vector and destination were incubated in a 1:1 ratio. _DH5 *E. coli* were transformed with 2_ μ_L of the reaction mix. Purified plasmid from an_ individual colony was sequence verified prior to use in the BACTH assay (see below).

Constructs for chlamydial transformation were created using the HiFi Cloning (New England Biolabs) protocol. Primers were designed to add a poly-Histidine (6xHis) tag to the gene of interest with the overlap to insert into the shuttle vector. Primers were generated using the NEBuilder^®^ assembly tool available from New England BioLabs (http://nebuilder.neb.com).

The backbone used was the pTLR2 derivative of the pASK plasmid (81). For the CRISPRi plasmid, the *S. aureus* dCas9 was PCR amplified from pX603-AAV-CMV::NLS-dSaCas9(D10A,N580A)-NLS-3xHA-bGHpA (a gift from Dr. F. Zhang; Addgene plasmid # 61594 (39)) and inserted into a derivative of pBOMB4-Tet::L2 (kind gift of Dr. T. Hackstadt, NIH; (82)) modified to weaken its ribosome binding site (Ouellette *in prep*). The gRNA cassettes were designed as previously described (38)), ordered as gBlock fragments from IDTDNA (Coralville, IA), and inserted into the BamHI site of the pBOMB4-Tet derivative encoding *Sa_dCas9* to produce, for example, the plasmid pBOMBLCRia::L2 (*clpP2X*). HiFi reactions were assembled according to the manufacturer’s protocol. The reaction was transformed into DH10ß *E. coli*, and isolated plasmid was verified by restriction enzyme digest and sequencing by Eurofins Genomics. Sequence verified plasmids were transformed into *dam-/dcm-E. coli* (New England BioLabs) to produce demethylated plasmid, which was verified as described earlier prior to transformation into *C. trachomatis* (see below).

For mutation of ClpX Walker B motif, Q5 mutagenesis (New England BioLabs) was used. Primers were designed encoding the E187A mutation for PCR linearization of the plasmid. ClpX BACTH constructs were used as a template for the PCR amplification, and plasmids were re-circularized by KLD reaction. The resulting reactions were transformed into DH5α*E. coli* for plasmid production. Plasmids were isolated, and mutations were verified by Sanger sequencing (Eurofins Genomics) prior to use in the BACTH system. These plasmids also served as template for the PCR reactions to produce PCR products for insertion of the mutant *clpX* gene into the pTLR2 plasmid.

Strains created or used in this study are listed in the supplementary material. Transformed *E. coli* strains were maintained on LB agar plates, with antibiotics as necessary. To extract chlamydial genomic DNA, EBs were subjected to heat and proteinase K treatment prior to phenol:chloroform extraction (83). Sodium hydroxide lysis was utilized for the extraction of *E. coli* genomic DNA. For cloning into the pLATE31 plasmid, the aLICator LIC Cloning and Expression Kit 3 (Thermo Scientific) was used according to the manufacturer’s specifications. Plasmids were first cloned into DH5α *E. coli* for plasmid propagation. Transformants were screened for inserts using colony PCR with Fermentas Master Mix (Thermo Scientific) and positive clones were grown for plasmid isolation (GeneJet Plasmid Miniprep Kit, Thermo Scientific). Sequence verified plasmids were then transformed into BL21(DE3) Δ*clpPAX E. coli* (46) for subsequent protein purification.

### Purification of Recombinant ClpX

His-tagged Ctr ClpX and Ctr ClpX_(E187A)_ were purified from 500 mL cultures of BL21(DE3) Δ *clppPAXE*. *coli* transformed with the respective plasmid based on the protocol described in (17). Samples were induced with 0.5 mM IPTG and incubated with shaking for 20 hours at 18°C. Cultures were pelleted and frozen at -80°C prior to purifications. Samples were suspended in buffer A (25 mM Tris Base [pH 7.5], 300 mM NaCl, and 10 mM Imidazole), sonicated, bound to HisPur Cobalt Resin (Thermo Scientific), and washed in buffer A Proteins were eluted from the resin using buffer B (25 mM Tris Base [pH 7.5], 300 mM NaCl, and 300 mM Imidazole). Buffer exchange for ATPase assay buffer (25 mM HEPES [pH 7.2], 200 mM KCl, 20 mM MgCl_2_, and 10% glycerol) was performed using a Millipore Amicon Ultra 15 filtration units (3 kDa cut-off). ClpX proteins were quantified using the Bio-Rad Protein assay, assessed for purity on 10% SDS-PAGE gels with Coomassie staining (Suppl. Fig. S9), and identified using anti-His-tag western blot. Blotting was performed using a mouse monoclonal anti-6x His antibody (1:1000; Millipore HIS.H8) and a goat anti-mouse IgG HRP conjugated secondary antibody (1:2000). Protein samples were aliquoted and stored at -80°C.

### *In Vitro* Analysis of ClpX Homo-Oligomerization

10 μg of purified protein was incubated at for 20 minutes at 37°C in oligomerization buffer (25 mM Tris Base [pH 7.5], 5 mM KCl, 5 mM MgCl_2_, 1 mM DTT, and 1% glycerol) prior to mixing with a 5x native sample buffer (5 mM Tris [pH 6.8], 38 mM glycine, 0.06% bromophenol blue). Assays were analyzed on a BioRad MiniProtean 4-20% gradient gel for Native-PAGE. Gels were assessed using Coomassie staining.

### Assessment of ClpX ATPase activity *in vitro*

A 49.5 µ L reaction containing 1.5 µ g of recombinant wild-type ClpX or ClpX_(E187A)_ in ATPase assay buffer (see above) supplemented with 1.7% DMSO was preincubated for 10 minutes at room temperature without ATP. Next, ATP dissolved in ATPase assay buffer was added to 1 µ M giving a final volume of 50 µ L, and the reaction was incubated at 30 C for 1.5 hours. After the 1.5 hours, the reactions were incubated for an additional 30 minutes at room temperature. 50 µ L of Kinase-Glo® reagent (Promega) was then added and incubated at room temperature for 10 minutes. Luminescence of the reaction, reflecting ATP not consumed by ClpX or ClpX_(E187A)_, was then measured using a BioTek Synergy H1 plate reader. Reactions were performed in duplicate at least three times with at least two independent protein preparations.

### Determining Protein-Protein Interactions with the BACTH System

The Bacterial Adenylate Cyclase Two-Hybrid (BACTH) assay was utilized to test interaction between ClpX wild-type and mutant (84). The genes of interest are translationally fused to one of either subunit, denoted as T18 and T25, of the *B. pertussis* adenylate cyclase toxin, which can complement adenylate cyclase deficient (Δ*cya*) DHT1 *E. coli*. Wild-type and mutant *clpX* genes cloned into one of the pST25, pSNT25, or pUT18C Gateway^®^ vectors was tested for both homotypic and heterotypic interactions (9, 80). Plasmids from each background were co-transformed into chemically competent DHT1 *E. coli*, which were plated on a double antibiotic minimal M63 medium selection plate supplemented with 0.5 mM IPTG for induction of the protein, 40 μg/mL Xgal, 0.04% casein hydrolysate, and 0.2% maltose. Leucine zipper motifs were used for controls in pKT25 and pUT18C backgrounds on the appropriate antibiotic selection plates because these have been previously shown to interact (85). Blue colonies, indicative of positive interaction, were screened using the β-galactosidase assay. Random positive colonies were selected and grown in M63 minimal media with the appropriate antibiotics. 0.1% SDS and chloroform were used to permeabilize the bacteria prior to addition of 0.1% o-nitrophenol-β-galactoside (ONPG). 1 M NaHCO_3_ was used to stop the reaction after precisely 20 minutes of incubation at room temperature. Absorbance at the 405 nm wavelength was recorded and normalized to bacterial growth (OD_600_), dilution factor, and time (in minutes) of incubation prior to stopping the reaction. Totals were reported in relative units (RU) of β-galactosidase activity.

### Chlamydial Transformation

The protocol followed was a modification of the method developed by Mueller and Fields (86) and as previously described (17). For transformation, 10^6^ *C. trachomatis* serovar L2 EBs (25667R) naturally lacking the endogenous plasmid were g of unmethylated plasmid in a volume of 50 μL CaCl_2_ at room temperature for 30 minutes. Reaction volume was sufficient for one well of a six well plate of McCoy mouse fibroblasts. Transformants were mixed with 1 mL of HBSS and added to 1 mL of HBSS in a six well plate. The plates were centrifuged at room temperature for 15 minutes, 400 xg. The plate was then incubated at 37° C for 15 minutes. After incubation, the HBSS was aspirated and replaced with antibiotic-free DMEM+10% FBS. 8 hours post-infection, the media was replaced with DMEM containing 1 μg/mL cycloheximide and 1 U/mL penicillin. Cells infected with transformants were passaged every 48 hours until a population of penicillin resistant bacteria was established. EBs were harvested and frozen in sucrose/phosphate (2SP; (64)) solution at -80° C.

### Determining the Effect of Overexpression of Wild-Type and Mutant Clp Proteins via Immunofluorescence and Inclusion Forming Unit Analysis

*C. trachomatis* transformants containing plasmids encoding the 6xHis-tagged protein of interest were used to infect a confluent monolayer of HEp2 cells. Penicillin treatment was maintained throughout the duration of the infection. At 10 hpi, samples were induced or not with 10 nM anhydrotetracycline (aTc). At the given timepoints, three wells of a 24 well plate were scraped in 2SP, vortexed with three 1 mm glass beads, and frozen at -80° C. At the same timepoint, a coverslip was fixed in 3.25% formaldehyde and 0.025% glutaraldehyde for two minutes, followed by permeabilization with cold 90% methanol for one minute. Coverslips were labeled with primary goat anti-major outer membrane protein (MOMP; Meridian, Cincinnati, OH), rabbit anti-6xHis (Abcam, Cambridge, MA), and DAPI. Appropriate donkey secondary antibodies were used (Invitrogen, Carlsbad, CA). Images were acquired on an Axio ImagerZ.2 equipped with Apotome.2 optical sectioning hardware and X-Cite Series 120PC illumination lamp. Frozen IFU samples were titrated onto a fresh monolayer of HEp2s without antibiotics. At 24 hpi, samples were fixed with methanol for 10 minutes, stained for MOMP, and enumerated.

### Genomic DNA Isolation and qPCR Enumeration of Genomic Equivalents

At 24 or 48 hpi, one well of a six well plate was scraped into the media overlay and pelleted at 17000 xg, 4° C for 15 minutes. Each sample was resuspended in 500 μL of cold PBS, frozen three times at -80° C, and processed using the Qiagen DNeasy Blood and Tissue Kit according to the manufacturer’s specifications. DNA concentrations were assessed using a spectrophotometer prior to dilution down to 5 ng/μL. 5 μL of the resulting dilution was used for a 25 μL qPCR reaction volume using SYBR^®^ Green PCR Master Mix (Applied Biosystems). Each reaction was performed in triplicate. A standard curve using Ctr L2 genomic DNA was generated for interpolation of sample Ct values. This experiment was performed three times for three biological replicates.

### Analysis of HctB Levels Upon Clp Overexpression

At 24 or 48 hpi, one well of a six well plate per test condition was rinsed twice with HBSS. To lyse the cells, 500 μL of denaturing lysis buffer (8 M Urea, 10 mM Tris, 2.5% 2-mercaptoethanol, 1% SDS) was added to each well and incubated for 15 minutes at room temperature. 300 units of Universal Nuclease (Pierce) per mL of lysis buffer was added immediately prior to addition to the wells. Following incubation, samples were centrifuged at 17000 xg, 4° C for 15 minutes to remove any insoluble material. Samples were quantitated using the EZQ Protein Quantitation Kit (Pierce). 50 μg of each sample was run in a 4-20% gradient SDS-PAGE gel (BioRad) and transferred to a PVDF 0.45 μm pore size membrane for 1 h at 300 mA. The membrane was probed using goat anti-MOMP (Meridian) and rabbit anti-HctB (generously provided by Dr. T. Hackstadt, NIH) primary antibodies followed by staining with donkey anti-goat 680 and donkey anti-rabbit 800 (LI-COR) secondary antibodies. The membrane was imaged on an Azure c600 imaging system. The channels were gray-scaled and equally contrast corrected, and the resulting images were used for integrated density measurement with FIJI software (87). To assess relative HctB levels, the HctB integrated density of each sample was normalized to its respective MOMP integrated density to avoid bias due to lower overall organism numbers. The ratios were then used to compare induced versus uninduced relative HctB levels. These experiments were performed three times for a total of three biological replicates.

### Transmission electron microscopy (TEM) assessment of the effect of overexpression of mutant Clp isoforms

Samples were infected and induced as previously discussed (see above). At 48 hpi, samples were fixed using 2% Glutaraldehyde, 2% Formaldehyde in 0.1M Sorensen’s phosphate buffer, pH 7.2. Samples were then stained post-fixation in 1% Osmium Tetroxide in water for 1 hour. Samples were dehydrated in an Ethanol series 50%, 70%, 90%, 95%, 100% 3 changes of 100%, all steps 15 minutes each and were then soaked in Propylene Oxide 100% 3 changes for 15 minutes each. Samples were left overnight in a fume hood in a 1:1 mixture of Propylene Oxide and Embed 812. The following day the samples were placed in molds with fresh Embed 812 and polymerized overnight in an oven set at 65° C. Blocks were thin sectioned 90 nanometers thick on a Leica UC6 Ultramicrotome using a Diatome diamond knife. Sections were placed on uncoated 200 mesh copper grids and stained with 2% Uranyl Acetate and Reynold’s Lead Citrate. Sections were examined on a FEI Tecnai G2 TEM operated at 80Kv.

### Confirmation of *clpP2X* and *incA* knockdown

Briefly, two wells of a six-well plate per condition were infected with pBOMBLCRia-*clpP2X* transformed CtrL2 at an MOI of 0.8. At either four or ten hpi, samples were or were not induced with 10 nM aTc. At each given timepoint, total RNA was collected using Trizol reagent (Invitrogen) and was extracted with chloroform as described previously (17, 83, 88–90). The aqueous layer was precipitated using isopropanol, as per the manufacturer’s instructions. Samples were DNase treated using the TURBO DNA-free kit (Ambion), and 1 µ g of the resulting RNA was reverse transcribed using SuperScript III reverse transcriptase (Invitrogen). Equal volumes of cDNA were loaded for each qPCR reaction. To extract genomic DNA, one well per condition was harvested and processed using the DNeasy blood and tissue kit (Qiagen) according to the manufacturer’s instructions as noted above. Samples were diluted to 5 ng/µ L, and 5 µ L of the resulting dilution was used per qPCR reaction. cDNA and gDNA samples were quantified using 25 µ L reactions with 2x SYBR PowerUP Green Master Mix (Invitrogen) analyzed on a QuantStudio 3 (Applied Biosystems) thermal cycler using the standard cycling conditions. A standard curve using purified wild-type CtrL2 genomic DNA was generated for sample quantification. Data are displayed as the ratio of cDNA to gDNA normalized to the 10h uninduced sample. For *incA* knockdown, HEp2 cells were infected with the pBOMBLCRia-*incA* transformant, induced with 10 nM aTc as above, and fixed at 24hpi with methanol. Cells were labeled with primary guinea pig anti-major outer membrane protein (MOMP; kind gift of Dr. E. Rucks, UNMC), rabbit anti-Sa_dCas9 (Abcam, Cambridge, MA), sheep anti-IncA (Dr. E. Rucks), and DAPI. Appropriate donkey secondary antibodies were used (Invitrogen). Coverslips were mounted on glass slides using ProLong Glass Antifade mounting media (ThermoFisher). Images were acquired on a Zeiss Axio ImagerZ.2 equipped with Apotome.2 optical sectioning hardware and X-Cite Series 120PC illumination lamp.

### Determination of the effect of *clpP2X* knockdown on Ctr

24-well plates of HEp2 cells were infected at an MOI of 0.8 with either pBOMBLCRia-*clpP2X* or pBOMBLCRia-*incA* transformed into CtrL2. Samples were induced or not at 4 hpi and were harvested, fixed, and titered as previously described. To monitor the expression of dCas9 and, if present, the complemented gene of interest, immunofluorescence samples on glass coverslips were fixed and permeabilized using ice cold 90% methanol at 24 or 48 hpi. Each sample was then stained using goat anti-MOMP (Meridian BioScience), rabbit anti-Sa_dCas9, and mouse anti-FLAG (Sigma-Aldrich) primary antibodies. Samples were then stained with the appropriate donkey secondary antibodies and were imaged as discussed above. Each titration was fixed using 4% formaldehyde and 0.025% glutaraldehyde to preserve GFP fluorescence. IFU counts of GFP positive inclusions are displayed as a percentage of the uninduced sample at the given timepoint. Plasmid retention for each condition is displayed as the percent of GFP positive to total number of inclusions for each condition.

### GFP_ssrA stability assay

Chlamydial transformants harboring either pBOMBmC-GFP(VAA) or pBOMBmC-GFP(VDD) plasmids were used to infect a fresh monolayer of McCoy mouse fibroblasts on coverslips in a 24 well plate. Samples were induced or not at 8 hpi. At 16 hpi, all media was replaced with fresh DMEM with or without the indicated compound. GFP expression was either continued through 20 hpi or was removed from 16 to 20 hpi for pulse/chase samples. At 20 hpi, all samples were formaldehyde/glutaraldehyde fixed as previously discussed. Coverslips were then mounted and imaged as previously discussed using a 40x objective. GFP and mCherry integrated densities were measured for approximately 60 fields of view on two separate coverslips per condition. These experiments were repeated for two biological replicates. To determine the ratio of GFP to mCherry, GFP integrated density measurements were divided by those of mCherry.

### Effect of ClpX-targeting compounds on chlamydial growth and host cell viability

Stocks of ClpX-specific inhibitor 334 and its derivative, 365, were synthesized as previously reported (44), resuspended at 25 mg/mL in DMSO, and frozen at -20°C. Methods for the synthesis, purification, and analysis of these compounds is available in Supplementary Information. A dose curve of treatment was performed to determine an inhibitory concentration of the compounds on Ctr, and 25 µ g/mL was chosen (data not shown). For the 24 and 48 h samples, 500 µ L of DMEM containing 25 µ g/mL of the compounds were added at 8 hpi, and samples were harvested at the indicated timepoint. For the time of infection samples, compounds were added at 15 minutes post-infection and removed at 8 hpi. For the reactivation samples, DMEM containing the respective compound was added at 8 hpi, washed out three times with HBSS at 24 hpi, and then replaced with DMEM only for 24 additional hours prior to harvest. To determine the effect on preformed EBs, compound was added at 24 hpi, and samples were harvested at 48 hpi. To harvest, three wells of a 24-well plate were scraped into 2SP, vortexed with 3 mm glass beads, and frozen at -80⁰ C. Samples were titrated onto a fresh monolayer of HEp2 cells with no treatment for enumeration. For assessment of cell viability, HEp2 cells were plate in 96 well dark wall plate and were treated as described for IFU experiments. All wells were then mixed with PrestoBlue HS reagent (Invitrogen), incubated for 10 minutes, and measured for fluorescence using a Tecan plate reader. Values are given as a percentage of the untreated sample.

## Acknowledgements

We thank Dr. H. Caldwell (NIH/NIAID) for eukaryotic cell lines, Dr. T. Hackstadt (RML/NIAID) for providing antibodies and the pBOMB4-Tet::L2 plasmid, Dr. P. Scott Hefty (KU) for the pTLR2-gfp::L2 plasmid, and Dr. Peter Sass (University of Tuebingen) for the BL21(DE3) Δ*clpPAX E. coli* strain used in this research. We thank Dr. Lisa Rucks antibodies used in this study and for critical review of the manuscript as well as members of the Ouellette and Rucks labs for helpful discussions of the work. This project was supported by a National Science Foundation CAREER award (1810599) and an NIAID/National Institutes of Health award (R21AI141933-01) to SPO and by University funds to DJF. This project was also funded by NIGMS/National Institutes of Health award to the Nebraska Center for Molecular Target Discovery and Development (1P20GM121316 01A1, PI: Robert Lewis, Project Leader, MCS).

---

**Figure S1: Transmission electron microscopy of the effects of overexpression of mutant Clp isoforms on chlamydial morphology. (A) Representative ClpP2_S98A_ uninduced or induced** samples at 48 hpi. Samples were induced or not with 10 nM aTc at 10 hpi and were fixed and processed at 48 hpi. Arrows indicate abnormal forms in the induced samples. Scale bar = 2 μm. **(B) Representative ClpX_E187A_ uninduced or induced** samples at 48 hpi. Samples treated, fixed, and processed as previously discussed. Arrows indicate abnormal forms with intrabacterial aggregates. Scale bar = 2 μm.

**Figure S2: (A) RT-qPCR of *clpP1*, *euo*, and *omcB* upon *clpP2X knockdown*.** Data shown are the average of three biological replicates, each with three technical replicates. Values are normalized to the 10h timepoint for each respective gene. Error bars represent standard deviation. Samples induced as indicated with 10 nM aTc. **(B) RT-qPCR of** *clpP2X* knockdown with *pcnB2* complementation. Values normalized to the ClpP2 10h samples.

**Figure S3: (A) Immunofluorescence staining** to confirm knockdown of IncA upon induction of dCas9 expression. Sa_dCas9 was induced or not at 4 hpi. Samples were harvested at 24 hpi and were stained for chlamydial MOMP (green), IncA (magenta), and DNA (Blue). Scale bar = 10 μm. **(B-C) Recoverable IFUs** and plasmid retention upon *incA* knockdown. **(D-E) Recoverable IFUs** and plasmid retention of *clpP2X* knockdown with *pcnB2* complementation.

**Figure S4: Representative immunofluorescence** images of **(A)** *clpP2X* knockdown with *pcnB2* complementation. Samples were induced as in IFU experiments. Chlamydial MOMP in green, FLAG (complemented gene) in red, dCas9 in magenta, and DNA in blue. **(B) Transcript levels** upon knockdown of *clpP2X* with *pcnB2* complementation. Data displayed are the values of three independent biological replicates of triplicate RT-qPCR reactions. Values are normalized to the 10h uninduced ClpP2 value for each experiment. Samples were induced using 10 nM aTc. **(C) IFU titer** following induction of knockdown and complementation at 4 hpi. Values are presented on a Log_10_ scale percent of the respective uninduced titer. **(D) Plasmid retention** of *clpP2X* knockdown with *pcnB2* complementation.

**Figure S5: Docking simulation of the ClpX inhibitor 334 on a ClpX model. (A) PDB structure** of 334. Inset is the 2D structure of the drug. **(B) Ribbon model** of docked 334 within the ClpX hexamer. The best scoring model is shown. Only the two ClpX subunits making contact with the model are shown (A in gray, B in seafoam green). The Walker A motif (red), Walker B motif (purple), sensor 1 motif (dark green), and sensor 2 motif (orange) of subunit A are colored for visualization. The arginine finger is labeled light green of subunit B. The center picture is oriented as outside of the complex looking inward, and the other images are rotations as indicated. **(C) Surface rendering** of the ClpX subunits with docked 334 are shown with coloration as in (B).

**Figure S6: Docking simulation of the ClpX inhibitor 365 on a ClpX model. (A) PDB structure** of 365. Inset is the 2D structure of the drug. **(B) Ribbon model** of docked 365 within the ClpX hexamer. The best scoring model is shown. Only the two ClpX subunits making contact with the model are shown (A in gray, B in seafoam green). The Walker A motif (red), Walker B motif (purple), sensor 1 motif (dark green), and sensor 2 motif (orange) of subunit A are colored for visualization. The arginine finger is labeled light green of subunit B. The center picture is oriented as outside of the complex looking inward, and the other images are rotations as indicated. **(C) Surface rendering** of the ClpX subunits with docked 365 are shown with coloration as in (B).

**Figure S7: Chemical disruption of ClpX function blocks degradation of an SsrA-tagged substrate in *C. trachomatis* L2.** Representative immunofluorescence images corresponding to data presented in Figure 6 of the main text. Representative images are of one field of view of one biological replicate. All samples were induced or not at 12 hpi with 20 nM aTc. Media was replaced with or without drug at 16 hpi, and aTc induction was maintained or removed (for pulse/chase samples). Samples were fixed at 20 hpi and were imaged as previously described using the same exposure time for all samples. Scale bars = 20 μm. Individual color channels were grayscaled and inverted to give clear distinction of fluorescent protein intensity.

**Figure S8: Chemical disruption of ClpX function is highly detrimental to *Chlamydia*. (A) Immunofluorescence assay (IFA)** of 24 hours post-infection (hpi). All drugs added at a final concentration of 25 μg/mL. Drugs were added at 8 hpi for the 24 h treatment samples. Drugs were added 15 minutes post-infection and removed at 8 hpi for the TOI: 8 h pulse samples. MOMP is stained in green, and DNA is stained with DAPI. Images were acquired on a Zeiss LSM800 microscope at 63x magnification. Scale bar = 10 μm. TOI = Time of Infection. **(B) IFA of 48 hpi** samples. Samples stained for MOMP (green) and DNA (blue). Scale bar = 10 μm. Drug was added at 8 hpi for 48 h and reactivation samples. Media removed, drug washed out, and media with drug added back to +Drug at 24 h and 48 h samples. Reactivation samples media replaced with DMEM, no drug. **(C) Recoverable inclusion forming units (IFUs)** from the indicated conditions. Totals present as Log_10_ IFUs recovered. Standard deviation displayed on graphs as error bars. **(D) PrestoBlue cell viability** assay upon drug treatment from 8 to 24 hpi. Values displayed as a % of the untreated samples.

**Figure S9: Example ClpX protein purifications.** Recombinant, 6x His-tagged ClpX/ClpX_(E187A)_ were purified using cobalt-based immobilized metal affinity chromatography. Samples were quantified and 1 and 5 µ g aliquots were run on 10% SDS-PAGE followed by staining with Coomassie brilliant blue. Three ClpX and two ClpX_(E187A)_ purification were performed using BL21(DE3) Δ*clpPAX E. coli*.

**Supplemental Table 1.** List of primers, plasmids, and strains used in this study.

## Notes

### Competing Interest Statement

The authors have declared no competing interest.

