## Supplementary figures and images for "The ClpX and ClpP2 Orthologs of *Chlamydia trachomatis* Perform Discrete and Essential Functions in Organism Growth and Development"

### Supplemental Figure S1

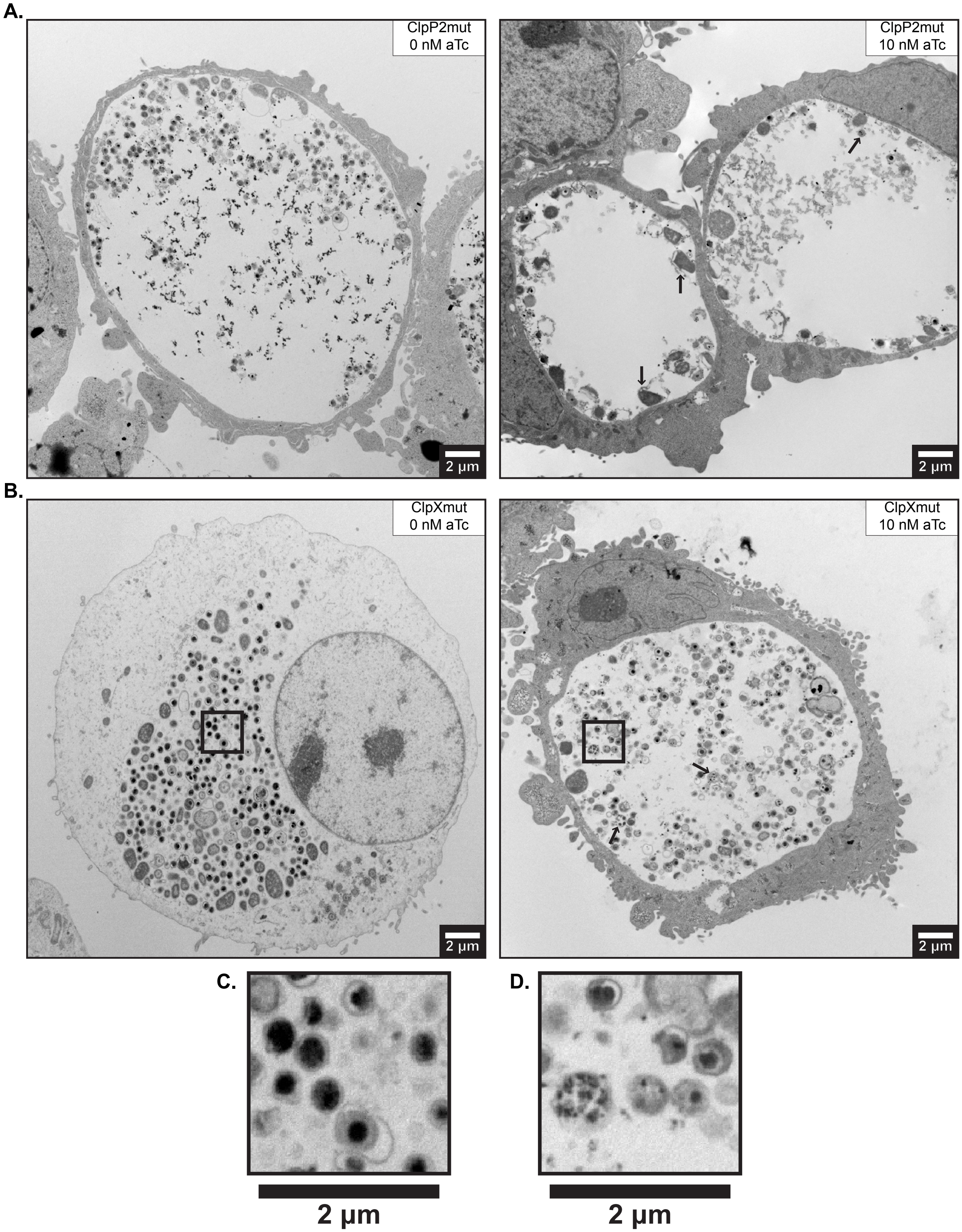

### Supplemental Figure S2

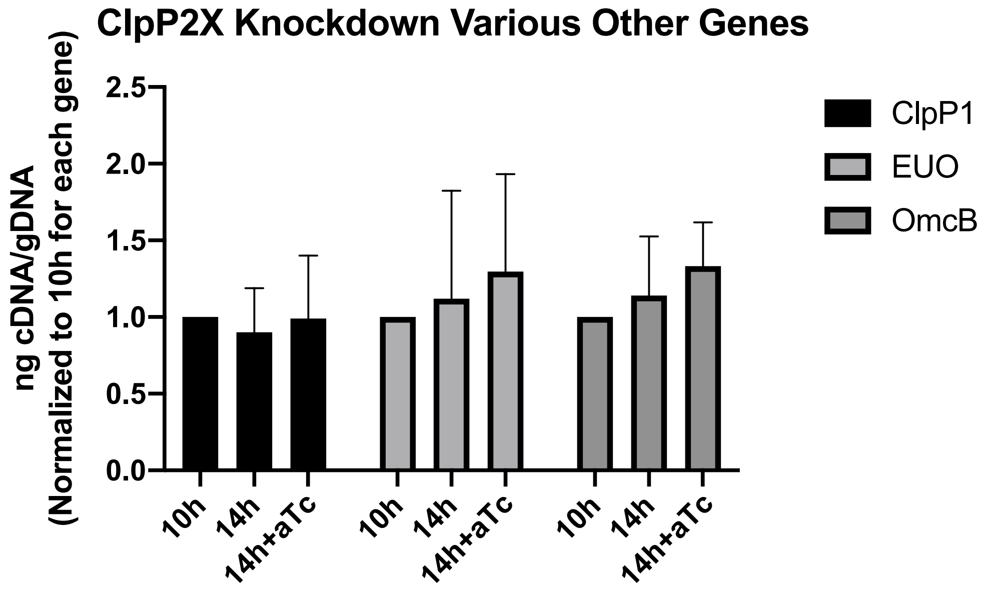

### Supplemental Figure S3

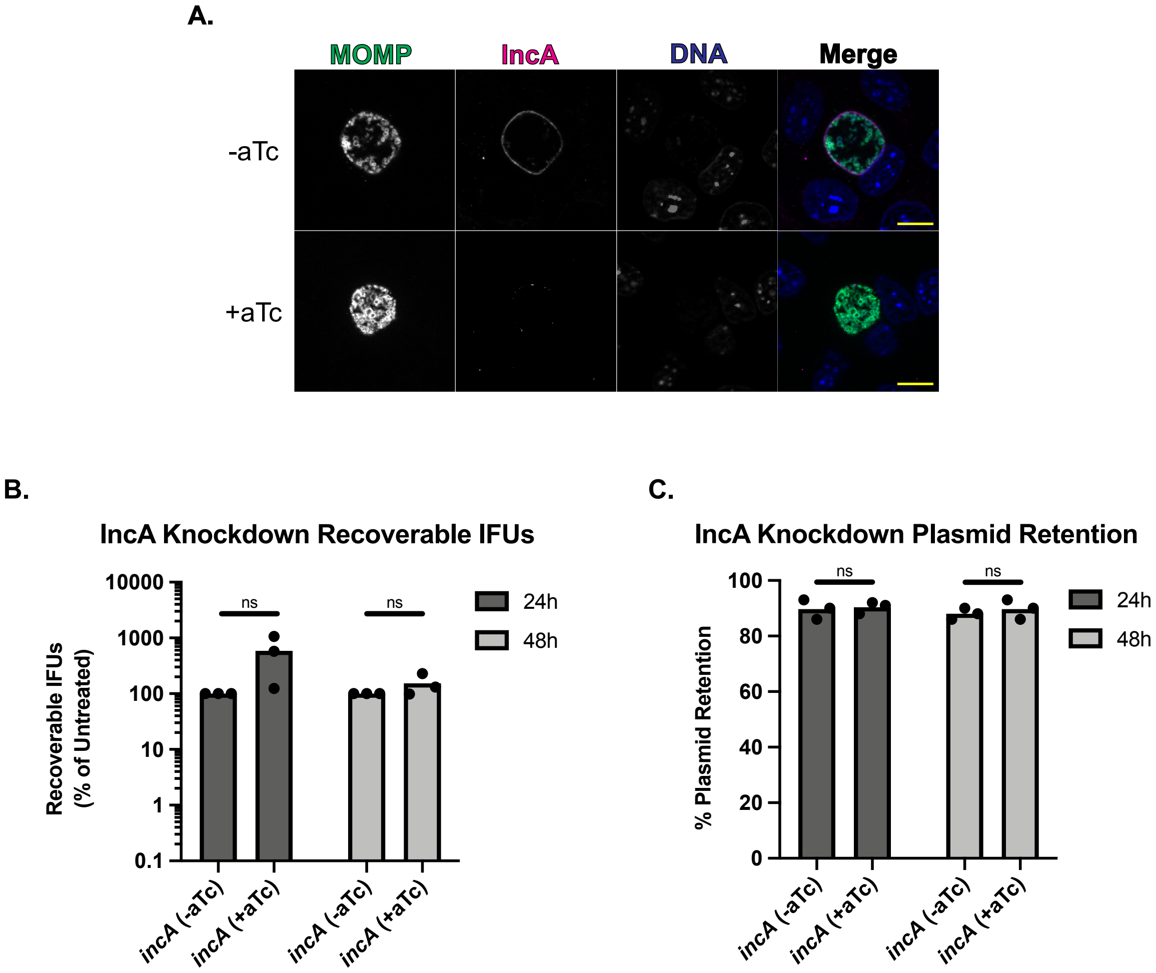

### Supplemental Figure S4

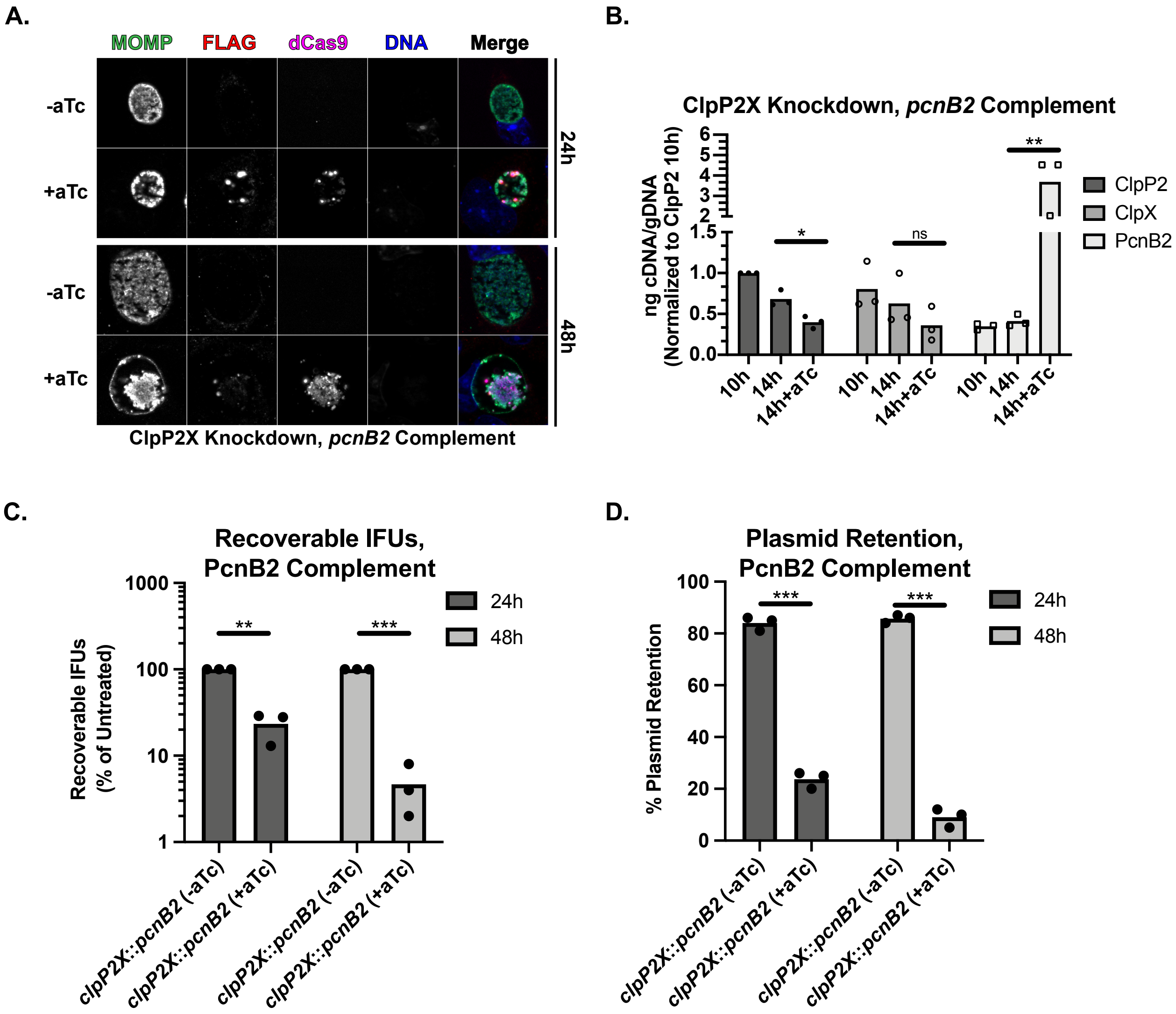

### Supplemental Figure S5

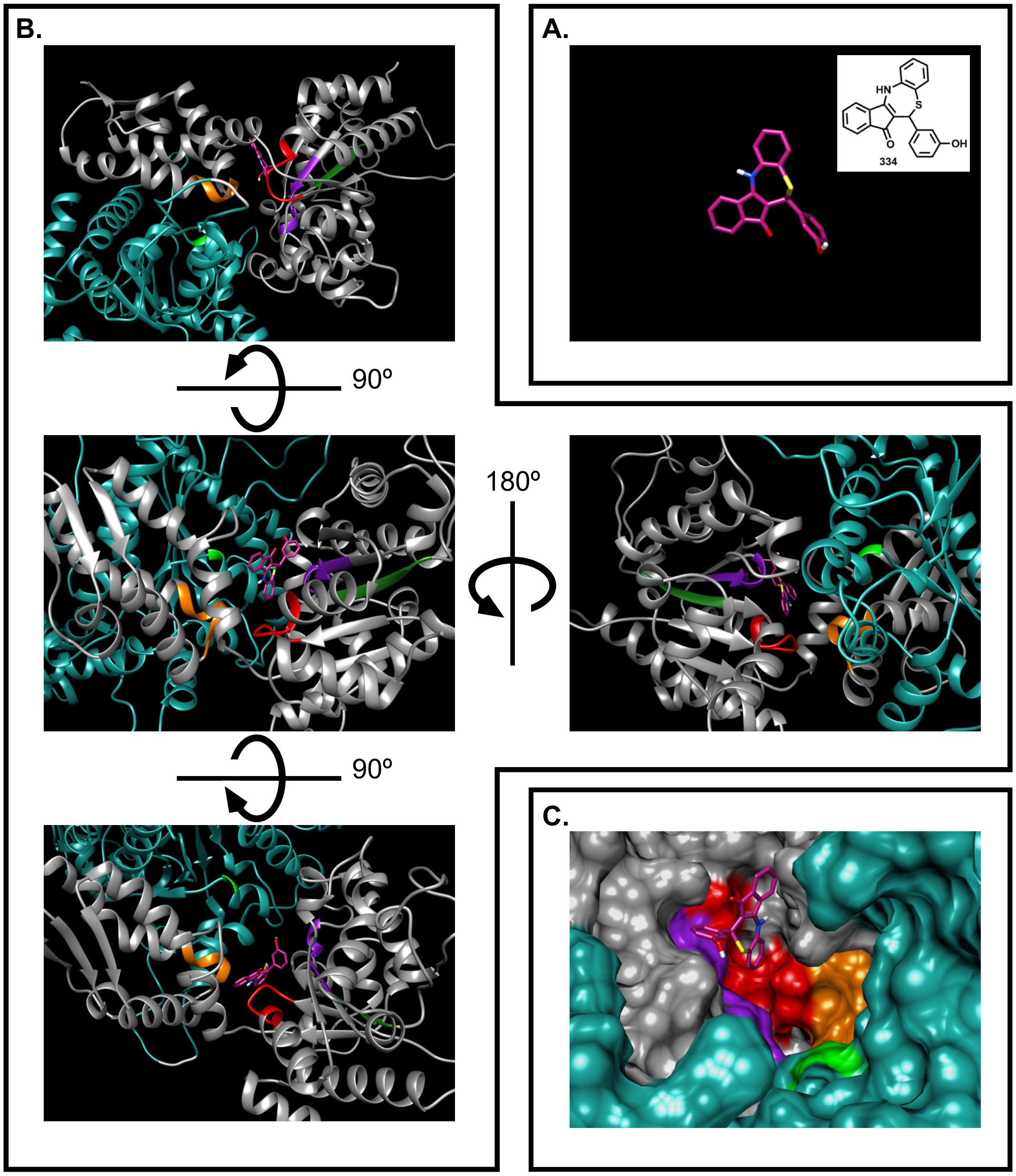


**Fig. S5**

### Supplemental Figure S6

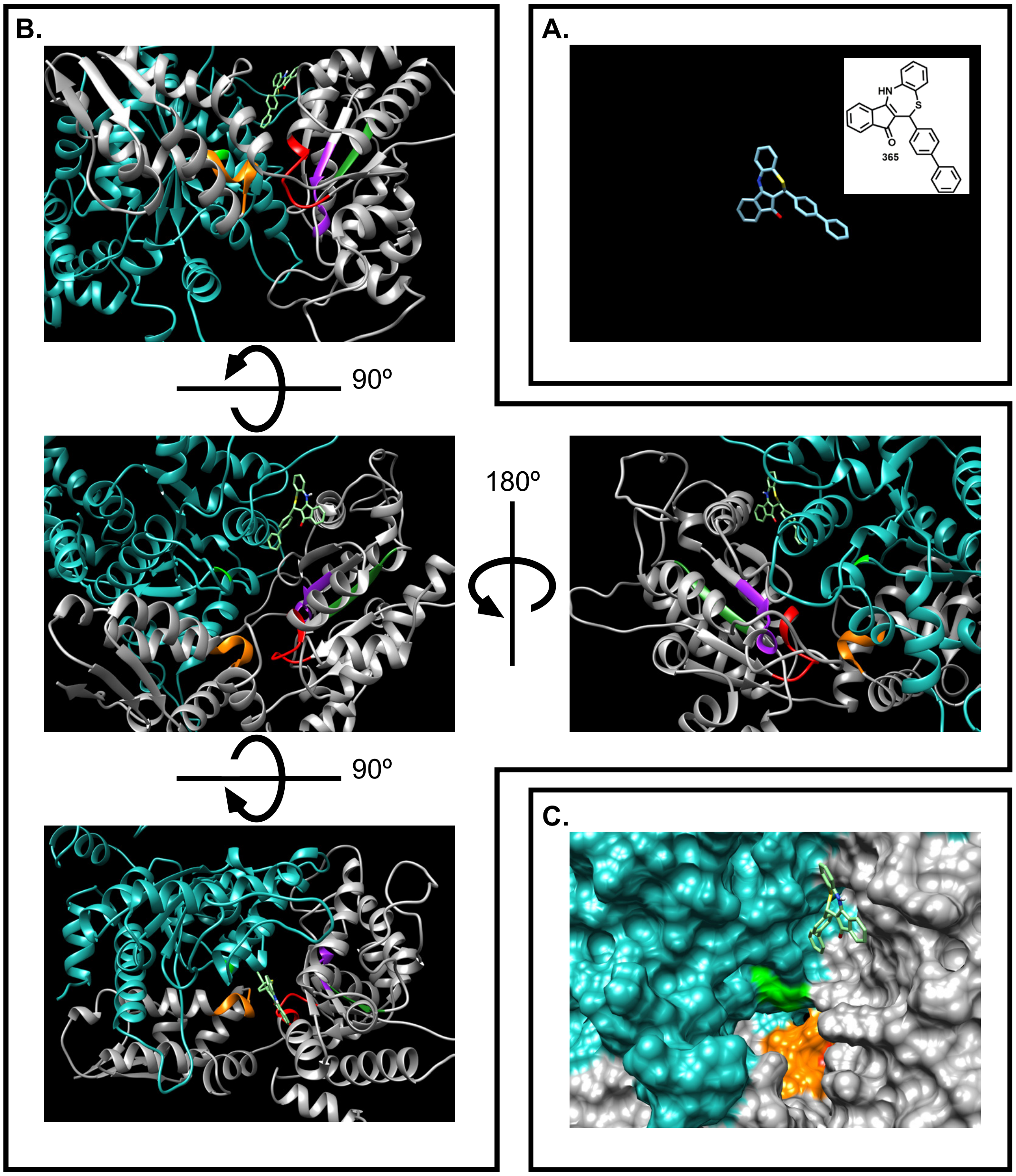


**Fig. S6**

### Supplemental Figure S7

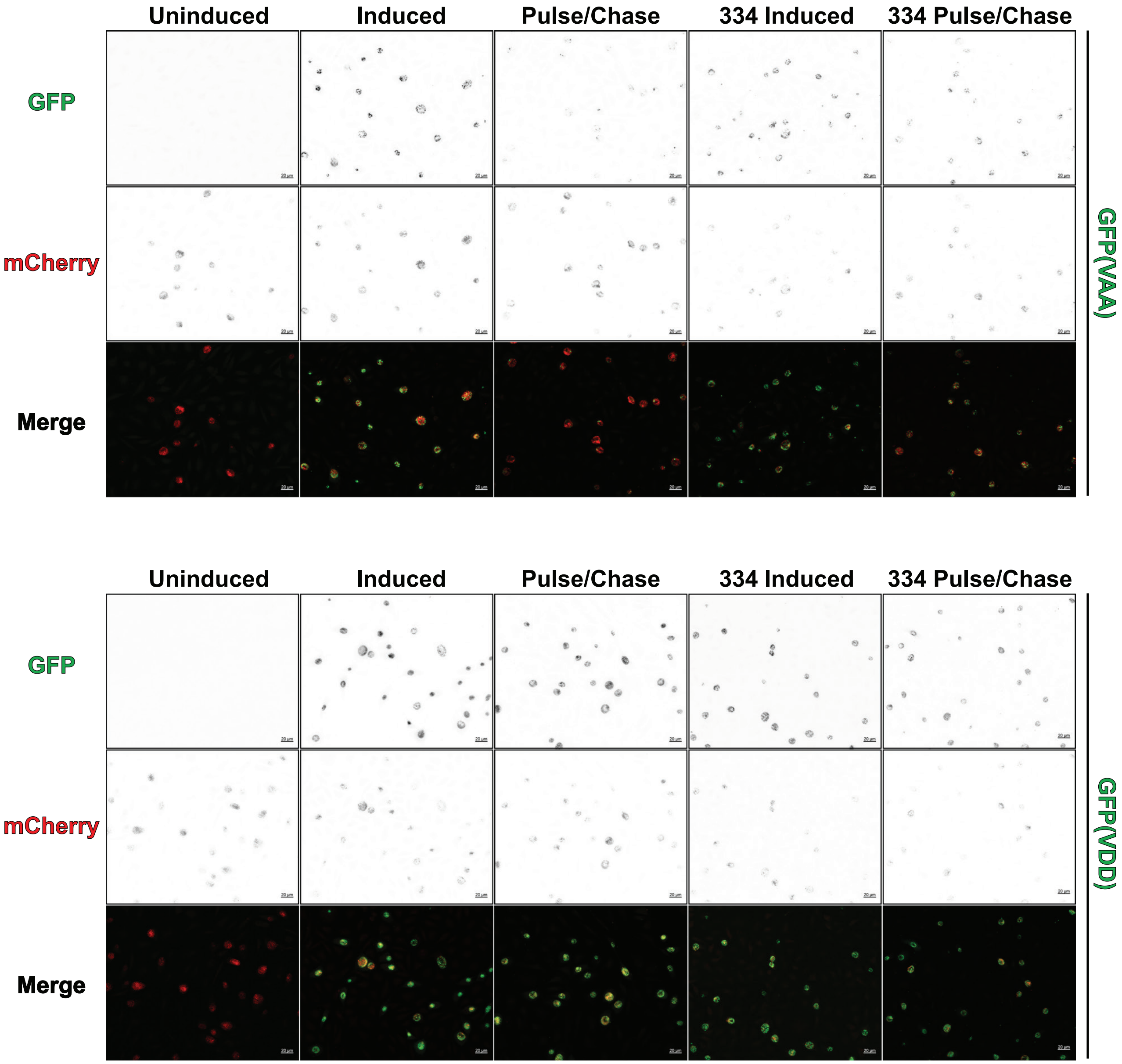

### Supplemental Figure S8

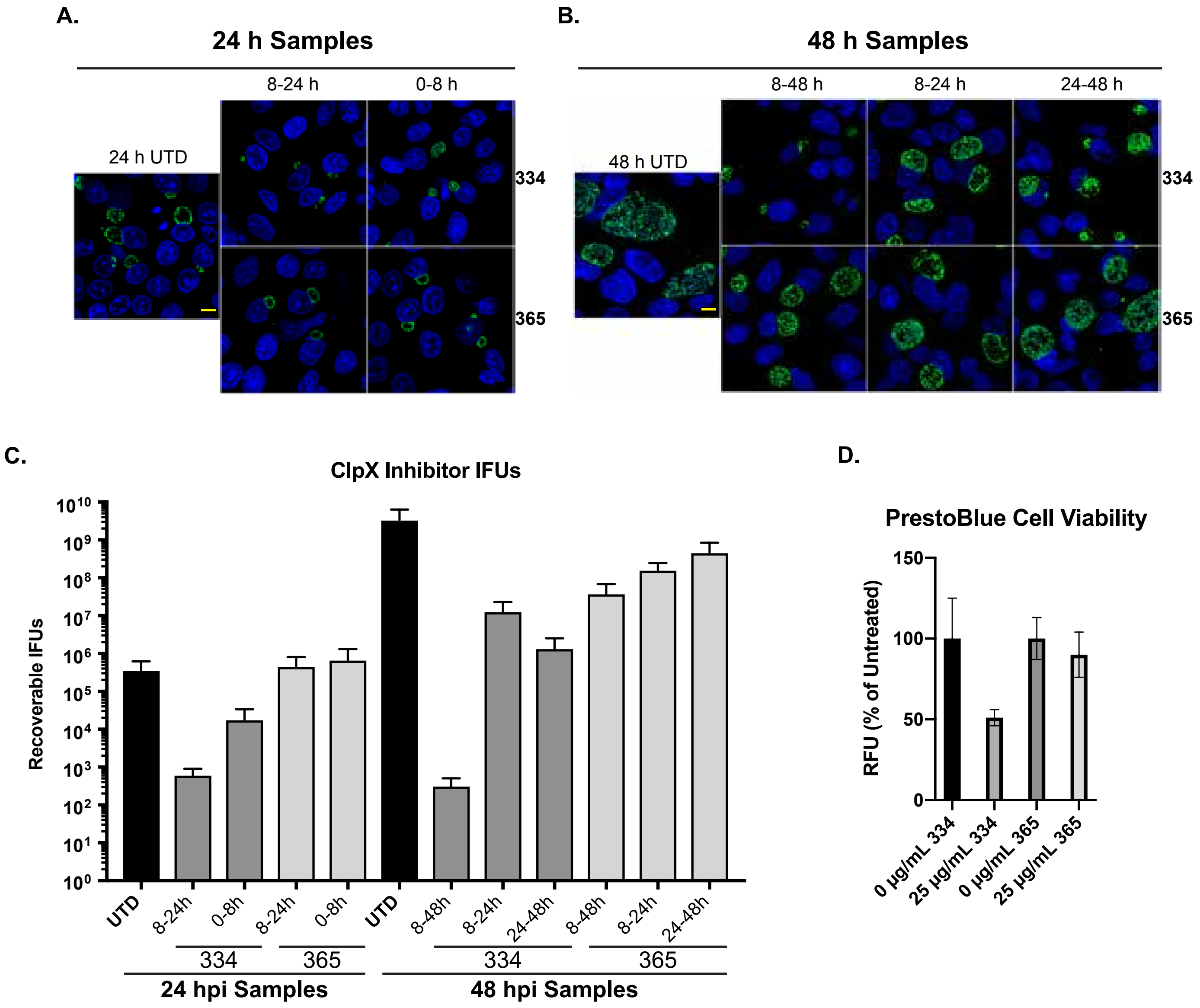

### Supplemental Figure S9

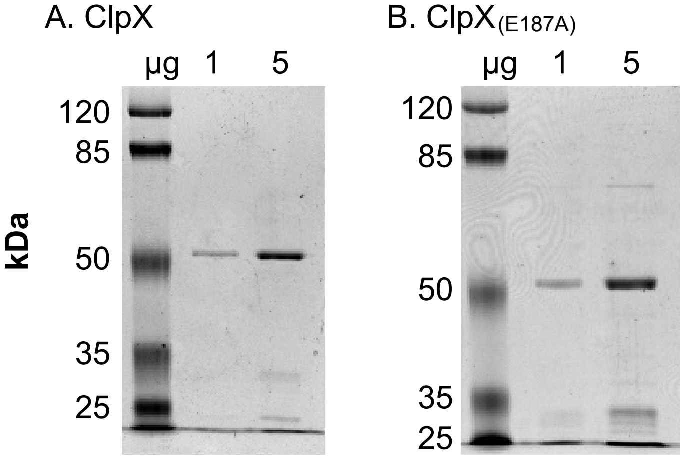


**Fig. S10**
