## Supplemental Table S1 for "The ClpX and ClpP2 Orthologs of *Chlamydia trachomatis* Perform Discrete and Essential Functions in Organism Growth and Development"

**Supplementary table 1. The list of Plasmids, Strains, and Primers**

| **Construct Plasmid** | **Relevant genotype** | **Ori** | **Source of Reference** |
| --- | --- | --- | --- |
| pTLR2-*clpP2* | *bla* P*tet*::*clpP2_*6xH | ColE1 | (1) |
| pTLR2-*clpP2_S98A_*_ | *bla* P*tet*:: *clpP2_S98A_*_6xH | ColE1 | (1) |
| pTLR2-*clpX* | *bla* P*tet*::*clpX_*6xH | ColE1 | This study |
| pTLR2-*clpX_E187A_* | *bla* P*tet*:: *clpX_E187A__*6xH | ColE1 | This study |
| pTLR2-*clpP2X* | *bla* P*tet*::*clpP2_*FLAG/*clpX*_6xH | ColE1 | This study |
| pTLR2-*clpP2_S98A_ X_E187A_* | *bla* P*tet*:: *clpP2_S98A__*FLAG/*clpX_E187A_*_6xH | ColE1 | This study |
| pST25-*clpX* | *aadA* P*_lac_*::*t25-attB1-clpX-attB2* | p15A | This study |
| pST25-*clpX_E187A_* | *aadA* P*_lac_*::*t25-attB1-clpX_E187A_-attB2* | p15A | This study |
| pUT18C-*clpX* | *bla* P*lac*::*t18-attB1-clpX-attB2* | ColE1 | This study |
| pUT18C-*clpX_E187A_* | *bla* P*lac*::*t18-attB1-clpX_E187A_-attB2* | ColE1 | This study |
| pKT25-*zip* | *aph* P*lac*::*t25-zip* | p15A | (2) |
| pUT18C-*zip* | *bla* P*lac*::*t18-zip* | ColE1 | (2) |
| pST25-DEST | *aadA* P*_lac_ attR1-[cat ccdB]-attR2* | p15A | (2) |
| pUT18C-DEST | *bla* P*_lac_ attR1-[cat ccdB]-attR2* | ColE1 | (2) |
| pDONR221 | *aph attP1-[cat ccdB]-attP2* | ColE1 | Invitrogen (Life Technologies) |
| pUT18C | *bla* P*lac*::*t18* | ColE1 | (2) |
| pENTR705 | *aph attL1-[clpX]-attL2* | ColE1 | This study |
| pLATE31-*clpX*_6xH | *bla* P*_tet_* P*_lac_*::*clpX*_6xH | pMB1 | This study |
| pLATE31-*clpX_E187A_*_6xH | *bla* P*_tet_* P*_lac_*::*clpX_E187A_*_6xH | pMB1 | This study |
| pBOMBLCRia(*clpP2X)*::L2 | *bla* P_dnaKm_::*clpP2X*_gRNA P*_tet_*::*Sa_dCas9vaa* | pUC19 | This study |
| pBOMBLCRia(*clpP2X)*-*clpP2_FLAG*::L2 | *bla* P_dnaKm_::*clpP2X*_gRNA P*_tet_*::*Sa_dCas9vaa/clpP2_FLAG* | pUC19 | This study |
| pBOMBLCRia(*clpP2X)*-*pcnB2_FLAG*::L2 | *bla* P_dnaKm_::*clpP2X*_gRNA P*_tet_*::*Sa_dCas9vaa/pcnB2_FLAG* | pUC19 | This study |
| pBOMBLCRia(*incA)*::L2 | *bla* P_dnaKm_::*incA*_gRNA P*_tet_*::*Sa_dCas9vaa* | pUC19 | Ouellette et al., *in prep* |
| pBOMB-G (no GFP) | *bla* P*_tet_*::mCherry | pUC19 | This study |
| pBOMBmC-*gfp(ssrA_VAA)* | *bla* P_Nm_::*mCherry* P*_tet_*::*gfp(ssrA_VAA)* | pUC19 | This study |
| pBOMBmC-*gfp(ssrA_VDD)* | *bla* P_Nm_::*mCherry* P*_tet_*::*gfp(ssrA_VDD)* | pUC19 | This study |

| ***E. coli* Strain** | **Relevant genotype** | **Source of Reference** |
| --- | --- | --- |
| DH5⍺ | *fhuA2 Δ(argF-lacZ)U169 phoA glnV44 Φ80 Δ(lacZ)M15 gyrA96 recA1 relA1 endA1 thi-1 hsdR17* | New England BioLabs |
| dam-/dcm- | *ara-14 leuB6 fhuA31 lacY1 tsx78 glnV44 galK2 galT22 mcrA dcm-6 hisG4 rfbD1 R(zgb210::Tn10) TetS endA1 rspL136 (*Str*^R^)dam13::Tn9 (*Cam*^R^) xylA-5 mtl-1 thi-1 mcrB1 hsdR2* | New England BioLabs |
| DHT1 | F^-^ *glnV44* (AS) *recA1 endA1 gyrA96* (Nal^R^) *thi-1 hsdR17 spoT1 rfbD1 cya-854 ilv-691 ::Tn10 (TetR)* | (3) |
| DH10β | *Δ(ara-leu) 7697 araD139 fhuA ΔlacX74 galK16 galE15 ϕ80dlacZΔM15 (e14-) recA1 relA1 endA1 nupG rpsL (*Str^R^*) rph spoT1 Δ(mrr-hsdRMS-mcrBC)* | New England BioLabs |
| *E*. *coli* dAPX-1 | *fhuA2, [lon], ompT, gal, [dcm], ΔhsdS, λ DE3 (λ sBamHIo ΔEcoRI-B int::(lacI::PlacUV5::T7 gene1) i21 Δnin5)* ΔclpX clpP::cam clpA::kan | (4) |

| **Primer name** | **Sequence** | **Features** | **Usage** |
| --- | --- | --- | --- |
| *attB*/*clpX*/ pDONR/5' | AATTAACAAGTTTGTACAAAAAAGCAGGCTTTATGACAAAAAAAAATCTTGCGGTCTGTTCTTTTTGT | For addition of 5’ *attB* recombination sites | for amplification *clpX* to use in Gateway Cloning |
| *attB*/*clpX*/ pDONR/3' | AATTACCACTTTGTACAAGAAAGCTGGGTTAGCAATCGCCTCTGGTGATTTCTGAATAATGACCG | For addition of 3’ *attB* recombination sites | for amplification *clpX* to use in Gateway Cloning |
| *clpX*/pTLR2/5' AgeI | CCCCCCCCCACCGGTATGACAAAAAAAAATCTTGCGGTCTGTTCT | 5’ end of *clpX* gene | for amplification of *clpX*_6xHis into pTLR2 |
| *clpX*/pTLR2/ 3'EagI 6xHis | ATATTCGGCCGTTAGTGATGGTGATGGTGATGAGCAATCGCCTCTGGTGATTTCTGAAT | Underline for 6xHis tag | for amplification of *clpX*_6xHis into pTLR2 |
| clp6xH/ (pTLR2)/LIC 3'v2 | ccatttttcacttcacaggtcaaccTTAGTGATGGTGATGGTGATG | lower case for plasmid overlap construction | for amplification of His-tagged *clp* genes into pTLR2 |
| *clpP2*/(pTLR2)/LIC 5' | tttgtttaactttaagaaggagataATGACGTTGGTACCATA | lower case for plasmid overlap construction | for amplification of *clpP2* into pTLR2 |
| *clpX*/(pTLR2)/ LIC 5' | tttgtttaactttaagaaggagataATGACAAAAAAAAATCTTGCG | lower case for plasmid overlap construction | for amplification of *clpX* into pTLR2 |
| *clpP2*/FLAG/ Rev | ACTTATCGTCGTCATCCTTGTAGTCAGACGCAATACTCTTATCTTTTG | Underline for FLAG tag | For addition of FLAG tag to *clpP2* |
| *clpP2X*/(pTLR2)/LIC P2 3' | taacaattctCTACTTATCGTCGTCATCCTTG | lower case for *clpP2X* IGR | for amplification of *clpP2*_FLAG into pTLR2 |
| *clpP2X*/(pTLR2)/LIC X 5' | tgacgacgataagtagagaattgttATGACAAAAAAAAATC | Underline for overlap cloning | for amplification of *clpX* into pTLR2 |
| *clpP2(FLAG)*/(pBOMBLCRia)/LIC Fwd | aaagggcgtagcagcataagTTTGTTTAACTTTAAGAAGGAGATAATG | lower case for plasmid overlap construction | For insertion of *clpP2* 3’ of *dCas9* in pLCRia |
| *clpP2(FLAG)*/(pBOMBLCRia)/LIC Rev | ttgaatggtcgaccggtacgCTACTTATCGTCGTCATC | lower case for plasmid overlap construction | For insertion of *clpP2* 3’ of *dCas9* in pLCRia |
| *pcnB2(FLAG)*/(pBOMBLCRia)/LIC Fwd | aaagggcgtagcagcataag*tttgtttaactttaagaaggagata*ATGACCCAGTCCACATTG | Italic for RBS | For insertion of *pcnB2* 3’ of *dCas9* in pLCRia |
| *pcnB2(FLAG)*/(pBOMBLCRia)/LIC Rev | ttgaatggtcgaccggtacg*ttacttatcgtcgtcatccttgtagtc*TTTCCAAAATCCCTTGCTC | Italic for addition of FLAG to *pcnB2* | For insertion of *pcnB2* 3’ of *dCas9* in pLCRia |
| P*_nm_* Fwd (for pBOMBmC) | cttttgctcacatggaattgGATGCCCGACGGTCTTTATAG | lower case for plasmid overlap construction | For P*_nm_*::*mCherry* insertion into pBOMB(-*gfp*) |
| P*_nm_* Rev (for pBOMBmC) | tagagaccatATCGGCTTCCTTTTGTAAATTTG | Underline for overlap with *mCherry* | For P*_nm_*::*mCherry* insertion into pBOMB-G |
| *mCherry* Fwd (for pBOMBmC) | ggaagccgatATGGTCTCTAAGGGCGAG | Underline for overlap with P*_nm_* | For P*_nm_*::*mCherry* insertion into pBOMB-G |
| *mCherry* Rev (for pBOMBmC) | agaaaaaacacctttaggcgTTATTTGTACAGCTCATCCATG | lower case for plasmid overlap construction | For P*_nm_*::*mCherry* insertion into pBOMB-G |
| *gfp*/pBOMBmC/Fwd | aaagaggagaaaggatctgcATGAGTAAAGGAGAAGCACTTTTC | lower case for plasmid overlap construction | For insertion of *gfp*(*ssrA*) into pBOMBmC |
| *gfp(VAA)*/pBOMBmC/Rev | tttgaatggtcgaccggtacTTAATCATCTACGCGTAGATC | lower case for plasmid overlap construction | For insertion of *gfp*(*ssrA_VAA*) into pBOMBmC |
| *gfp(VDD)*/pBOMBmC/Rev | tttgaatggtcgaccggtacTTAAGCAGCTACGCGTAG | lower case for plasmid overlap construction | For insertion of *gfp*(*ssrA_VDD*) into pBOMBmC |
| *clpX*/mutE187A/Q5_F | TTACATTGAT**GCA**ATCGATAAAATTGGTCG | Underline for E187A mutation | for mutation of the *clpX* Walker B motif |
| *clpX*/mutE187A/Q5_R | ATAATGCCTCGCTCTGCTC | For E187A mutagenesis | for mutation of the *clpX* Walker B motif |
| ct443 *omcB* | CGGTAGGATCTCCCTATCCTATT | Forward qPCR primer | For qPCR of *omcB* |
| ct443 *omcB* | CGAACTCTGCTTCACATGGTA | Reverse qPCR primer | For qPCR of *omcB* |
| ct446 *euo* | CGAAGACTACTCGTTGGGAAATA | Forward qPCR primer | For qPCR of *euo* |
| ct446 *euo* | AACAGAAGCTCTCCTTGATAAGT | Reverse qPCR primer | For qPCR of *euo* |
| ct441 *clpP_1* | GATGCTGGGTTTGCTGTTTG | Forward qPCR primer | For qPCR of *clpP1* |
| ct441 *clpP_1* | CAGATCCCATAGATGCTGCTAAA | Reverse qPCR primer | For qPCR of *clpP1* |
| ct706 *clpP_2* | GTTAGCGATTTACGACACCATTC | Forward qPCR primer | For qPCR of *clpP2* |
| ct706 *clpP_2* | CCCTTTGTCCCTGCAGATAATA | Reverse qPCR primer | For qPCR of *clpP2* |
| ct705 *clpX* | GGTTGCCGTCTATAACCACTATAA | Forward qPCR primer | For qPCR of *clpX* |
| ct705 *clpX* | AGATCCTGTTGGGCCTAGTA | Reverse qPCR primer | For qPCR of *clpX* |
| ct704 *pcnB2* | GACGATGCTATTCGTAGGGATTT | Forward qPCR primer | For qPCR of *pcnB2* |
| ct704 *pcnB2* | TCTTTCTAAGTCGGCTCTACCT | Reverse qPCR primer | For qPCR of *pcnB2* |

| **gBlock Name** | **Sequence** | **Features** | **Usage** |
| --- | --- | --- | --- |
| *incA*_IGR gRNA gBlock | *tgtgaaagtgggtcttaagacgtcg*gtactgcatgtgacgcacgtagatcatgcaTTCACCGGTGGAGACGGTTTTCTTATAATGACACC**AATTTTTATCATATAAAGCCC**GTTTTAGTACTCTGGAAACAGAATCTACTAAAACAAGGCAAAATGCCGTGTTTATCTCGTCAACTTGTTGGCGAGATTTTTCAAATAAAACGAAAGGCTCAGTCGAAAGACTGGGCCTTTCGTTTTATcaacagcggtctactgaatctgagctagtg*cgtgatataattaaaattatattca* | Italicized sequence represents pBOMB flanking regions for HiFi insertion.  Lowercase sequence is random spacer.  Bolded sequence is the target gRNA. | For CRISPRi knockdown of *incA* |
| *clpP2X*_IGR gRNA gBlock | *tgtgaaagtgggtcttaagacgtcg*gtactgcatgtgacgcacgtagatcatgcaTTCACCGGTGGAGACGGTTTTCTTATAATGACACC**ATACACCTACGCATCAAAATG**GTTTTAGTACTCTGGAAACAGAATCTACTAAAACAAGGCAAAATGCCGTGTTTATCTCGTCAACTTGTTGGCGAGATTTTTCAAATAAAACGAAAGGCTCAGTCGAAAGACTGGGCCTTTCGTTTTATcaacagcggtctactgaatctgagctagtg*cgtgatataattaaaattatattca* | Italicized sequence represents pBOMB flanking regions for HiFi insertion.  Lowercase sequence is random spacer.  Bolded sequence is the target gRNA. | For CRISPRi knockdown of *clpP2X* |
| *gfp(VDD)* gBlock | tccaggggcccctgggatccAGTAAAGGAGAAGCACTTTTCACTGGAGTTGTCCCAATTCTTGTTGAATTAGATGGTGATGTTAATGGGCACAAATTTTCTGTCAGTGGAGAGGGTGAAGGTGATGCAACATACGGAAAACTTACCCTTAAATTTATTTGCACTACTGGAAAACTACCTGTTCCATGGCCAACACTTGTCACTACTCTTACGTATGGTGTTCAATGCTTTTCAAGATACCCAGATCATATGAAACGGCATGACTTTTTCAAGAGTGCCATGCCCGAAGGTTATGTACAGGAAAGAACTATATTTTTCAAAGATGACGGGAACTACAAGACACGTGCTGAAGTCAAGTTTGAAGGTGATACCCTTGTTAATAGAATCGAGTTAAAAGGTATTGATTTTAAAGAAGATGGAAACATTCTTGGACACAAATTGGAATACAACTATAACTCACACAATGTATACATCATGGCAGACAAACAAAAGAATGGAATCAAAGTTAACTTCAAAATTAGACACAACATTGAAGATGGAAGCGTTCAACTAGCAGACCATTATCAACAAAATACTCCAATTGGCGATGGCCCTGTCCTTTTACCAGACAACCATTACCTGTCCACACAATCTGCCCTTTCGAAAGATCCCAACGAAAAGAGAGACCACATGGTCCTTCTTGAGTTTGTAACAGCTGCTGGGATTACACATGGCATGGATGAACTATACAAGGCCGAACCGAAGGCTGAATGCGAAATTATCAGCTTCGCTGATCTCGAAGATCTACGC**GTAGATGAT**TAAgtcgactcgagcggccgcatcgtgactgac | Lowercase represents non-specific sequence. Uppercase represents *gfp* sequence. Bolded letters indicate *VDD* of *ssrA* tag. | Template for amplification of *gfp(VDD)* for insertion into pBOMBmC |
| *gfp(VAA)* gBlock | tccaggggcccctgggatccAGTAAAGGAGAAGCACTTTTCACTGGAGTTGTCCCAATTCTTGTTGAATTAGATGGTGATGTTAATGGGCACAAATTTTCTGTCAGTGGAGAGGGTGAAGGTGATGCAACATACGGAAAACTTACCCTTAAATTTATTTGCACTACTGGAAAACTACCTGTTCCATGGCCAACACTTGTCACTACTCTTACGTATGGTGTTCAATGCTTTTCAAGATACCCAGATCATATGAAACGGCATGACTTTTTCAAGAGTGCCATGCCCGAAGGTTATGTACAGGAAAGAACTATATTTTTCAAAGATGACGGGAACTACAAGACACGTGCTGAAGTCAAGTTTGAAGGTGATACCCTTGTTAATAGAATCGAGTTAAAAGGTATTGATTTTAAAGAAGATGGAAACATTCTTGGACACAAATTGGAATACAACTATAACTCACACAATGTATACATCATGGCAGACAAACAAAAGAATGGAATCAAAGTTAACTTCAAAATTAGACACAACATTGAAGATGGAAGCGTTCAACTAGCAGACCATTATCAACAAAATACTCCAATTGGCGATGGCCCTGTCCTTTTACCAGACAACCATTACCTGTCCACACAATCTGCCCTTTCGAAAGATCCCAACGAAAAGAGAGACCACATGGTCCTTCTTGAGTTTGTAACAGCTGCTGGGATTACACATGGCATGGATGAACTATACAAGGCCGAACCGAAGGCTGAATGCGAAATTATCAGCTTCGCTGATCTCGAAGATCTACGC**GTAGCTGCT**TAAgtcgactcgagcggccgcatcgtgactgac | Lowercase represents non-specific sequence. Uppercase represents *gfp* sequence. Bolded letters indicate *VAA* of *ssrA* tag. | Template for amplification of *gfp(VAA)* for insertion into pBOMBmC |

1. Wood NA, Chung K, Blocker A, Rodrigues de Almeida N, Conda-Sheridan M, Fisher DJ, Ouellette SP. 2018. Initial Characterization of the Two ClpP Paralogs of *Chlamydia trachomatis* Suggests Unique Functionality for Each. *Journal of Bacteriology*.
2. Karimova G, Pidoux J, Ullmann A, Ladant D. 1998. A bacterial two-hybrid system based on a reconstituted signal transduction pathway. *Proceedings of the National Academy of Sciences of the United States of America.* 95:5752-5756.
3. Dautin N, Karimova G, Ullmann A, Ladant D. 2000. Sensitive genetic screen for protease activity based on a cyclic AMP signaling cascade in *Escherichia coli*. *Journal of bacteriology*. 182:7060-7066.
4. Pan S, Malik IT, Thomy D, Henrichfreise B, Sass P. 2019. The functional ClpXP protease of *Chlamydia trachomatis* requires distinct clpP genes from separate genetic loci. *Scientific Reports* 9:14129.
